# Spatiotemporal whole-brain dynamics of auditory patterns recognition

**DOI:** 10.1101/2020.06.23.165191

**Authors:** L. Bonetti, E. Brattico, F. Carlomagno, J. Cabral, A. Stevner, G. Deco, P.C. Whybrow, M. Pearce, D. Pantazis, P. Vuust, M.L Kringelbach

## Abstract

Music is a non-verbal human language, built on logical structures and articulated in balanced hierarchies between sounds, offering excellent opportunities to explore how the brain creates meaning for complex spatiotemporal auditory patterns. Using the high temporal resolution of magnetoencephalography in 70 participants, we investigated their unfolding brain dynamics during the recognition of previously memorized J.S. Bach’s musical patterns from prelude in C minor BWV 847 compared to novel patterns matched in terms of entropy and information content. Remarkably, the recognition of the memorized music ignited a widespread brain network comprising primary auditory cortex, superior temporal gyrus, insula, frontal operculum, cingulate gyrus, orbitofrontal cortex, basal ganglia, thalamus and hippocampus. Furthermore, measures of both brain activity and functional connectivity presented an overall increase over time, following the evolution and unfolding of the memorized musical patterns. Specifically, while the auditory cortex responded mainly to the first tones of the patterns, the activity and synchronization of higher-order brain areas such as cingulate, frontal operculum, hippocampus and orbitofrontal cortex largely increased over time, arguably representing the key whole-brain mechanisms for conscious recognition of auditory patterns as predicted by the global neuronal workspace hypothesis. In conclusion, our study described the fine-grained whole-brain activity and functional connectivity dynamics responsible for processing and recognition of previously memorized music. Further, the study highlights how the use of musical patterns in combination with a wide array of analytical tools and neuroscientific measures spanning from decoding to fast neural phase synchronization can shed new light on meaningful, complex cognitive processes.

## Introduction

Higher human brain function crucially depends on advanced pattern recognition and prediction of complex information [1, 2]. This fast processing of information available in the present moment and its relation to the past and future is not effortless but demanding [3, 4]. Specifically, this processing depends on a complex chain of events, including stimulus identification, extraction of meaningful features, comparison with memory representation and prediction of upcoming stimuli [5, 6].

For instance, a sequence of images, sounds, or words can be encoded and stored at several levels of detail, from specific items and their timing to abstract structure. This phenomenon is thought to occur through a process of strengthening the synapses that form the neural circuit responsible for the encoding of the information. Then, reengaging successfully the neural circuit formed during the encoding phase is crucial to properly predict and recognize the information conveyed by the constantly incoming external stimuli [7]. Neuroscientific research has widely investigated this central topic, focusing especially on processing and recognition of images and words [8, 9]. Additionally, in the auditory domain, patterns have been extensively studied in the context of prediction error and automatic detection of environmental irregularities, as indexed by event-related components such as N100 and mismatch negativity (MMN) [10, 11]. Indeed, this literature has established that the brain is remarkably capable of recognizing patterns and detecting deviations, which are key capacities to accomplish any complex cognitive function.

However, even if the interest in pattern recognition has increased across scientific fields and neuroscience, the brain mechanisms responsible for conscious recognition of patterns acquiring meaning over time are not fully understood yet. Indeed, a recent paper by Stanislas Dehaene and colleagues [12] explicitly pointed out that future neuroscientific studies are called for to reveal the role of different brain areas in extracting and retrieving temporal sequences.

Thus, in this study, we focused on music, the human art that mainly acquires meaning through the combination of its constituent elements extended over time [13, 14]. Music is an excellent tool for investigating the brain, which similar to language has been proposed to be a universal of human culture [15]. Indeed, beyond the strong emotional content evoked, music contains complex logical structures, articulated in carefully balanced hierarchies between sounds [14, 16], yielding to meaningful messages and information that can be processed and recognized. Therefore, music is a great tool for investigating higher human brain processes implicated in the recognition of patterns unfolding over time.

Here, we selected the prelude in C minor BWV 847 composed by Johann Sebastian Bach, given its repetitive and memorable structure [17, 18], which makes it highly suitable for the current neuroscientific investigation. We extracted short musical patterns from the prelude and created novel melodies matched in terms of entropy, information content and main acoustic features. This procedure, combined with the use of state-of-the-art neuroscientific techniques such as magnetoencephalography (MEG) and magnetic resonance imaging (MRI), allowed us to investigate the fundamental brain mechanisms underlying the conscious recognition of auditory patterns evolving over time.

## Results

### Experimental design and data analysis overview

In this study, we wanted to characterize the fine-grained spatiotemporal dynamics of brain activity and connectivity during recognition of previously memorized auditory patterns. During a session of magnetoencephalography (MEG), 70 participants listened to a musical instrumental digital interface (MIDI) version of the full prelude in C minor BWV 847 composed by Bach and tried to memorize it as much as possible. As described in the Methods and depicted in **Figure 1A** and **Figure SF1**, participants were then presented with short musical melodies corresponding to excerpts of the Bach’s prelude and carefully matched novel musical sequences (in terms of information content, entropy, and main acoustic features, see Methods for details). Participants were asked to indicate whether each musical excerpt was extracted from the Bach’s prelude or was a novel melodic pattern.

**Figure 1.**
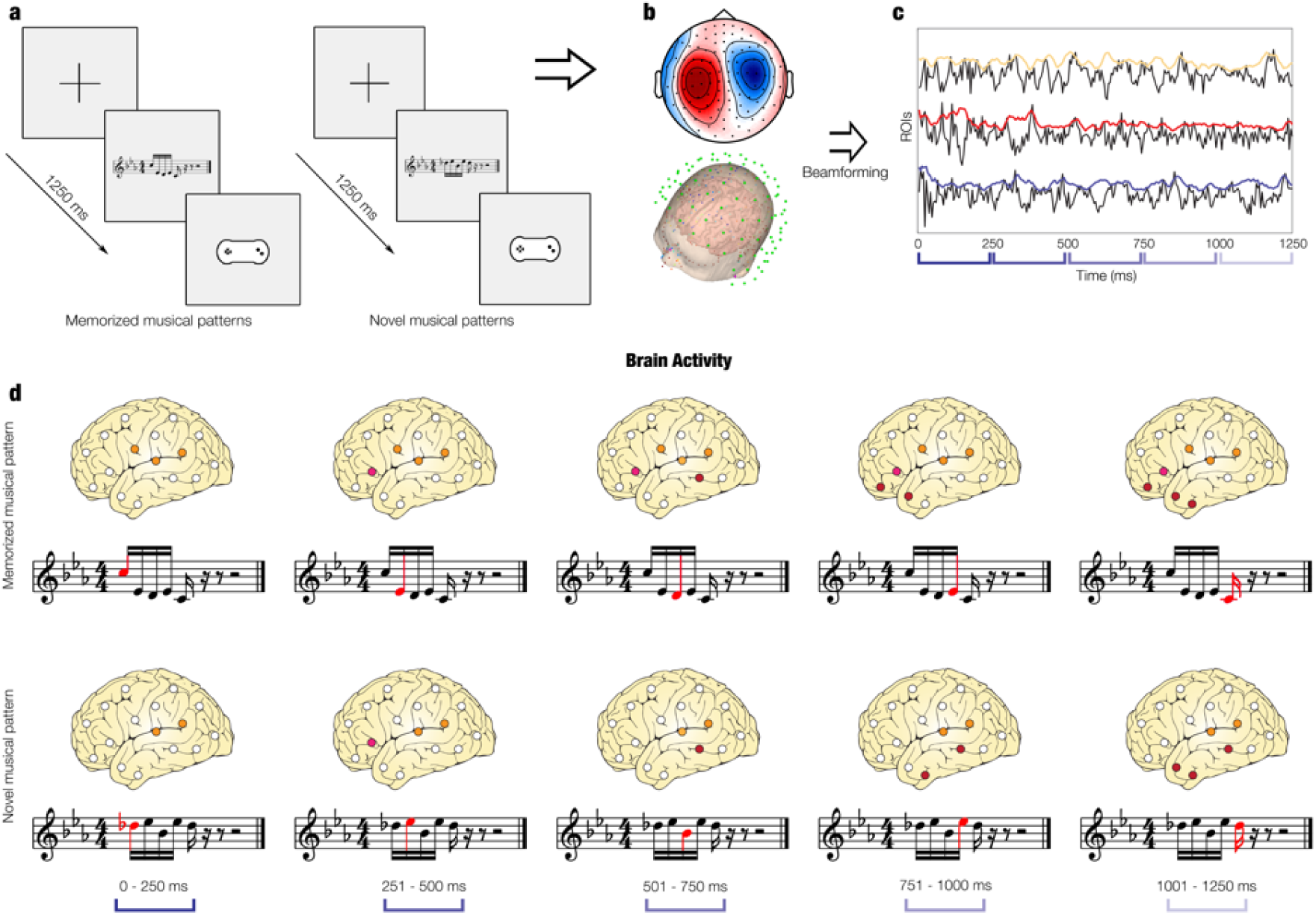
Experimental design and analysis methods. **a -** Graphical schema of the old/new paradigm. One at a time, several five-tone musical patterns (melodies) were presented. They could either be taken from the whole Bach’s prelude that participants previously listened to (memorized musical pattern, ‘old’) or could be novel musical patterns (‘new’). In this figure, we depicted at first an example of a memorized musical pattern (‘old’) pattern (left, 2^nd^ square) with the relative response pad that participants used to state whether they recognized the excerpt as ‘old’ or ‘new’ (left, 3^th^ square). Then, we depicted an example of novel musical pattern (‘new’) (right, 2^nd^ square). The total number of trials was 80 (40 memorized and 40 novel musical patterns) and their order was randomized. **b –** We collected, pre-processed and analyzed MEG sensor data by employing multivariate pattern analysis and MCS on univariate tests. **c –** We beamformed MEG sensor data into source space, providing time series of activity originating from brain locations. **d –** We studied the source brain activity underlying the processing of each tone of the musical sequences for both experimental conditions.

The analysis pipeline used in this study is illustrated in **Figure 1B** (and described in detail in the Methods). This focused on extracting results using four main measures of brain functioning: 1) MEG sensor space activity, 2) beamformed source localized activity, 3) static source localized connectivity and 4) dynamic source localized connectivity.

First, we used multivariate pattern analysis and Monte Carlo simulations (MCS) on univariate tests of MEG sensor data. Second, we were interested in finding the brain sources of the observed differences and therefore we reconstructed the sources of the signal using a beamforming algorithm (**Figure 1C**) to track the brain activity related to each tone of the musical sequences (**Figure 1D**). Third, complementing our brain activity results, we computed the functional connectivity between different brain regions. We calculated the static functional connectivity by computing Pearson’s correlations between the envelopes of each pair of brain areas. Fourth, we investigated the functional connectivity dynamics.

**Figure 2A** depicts the Hilbert transform used to estimate the instantaneous phase synchronization between brain areas, while **Figure 2B** and **2C** illustrate the instantaneous functional connectivity matrices showing the connectivity patterns characterizing the brain at each time point. Similar to our analysis of brain activity, we analyzed the brain connectivity patterns related to each tone of the musical sequences, as shown in **Figure 2D**. Here, we assessed both whole-brain connectivity and degree centrality of brain regions.

**Figure 2.**
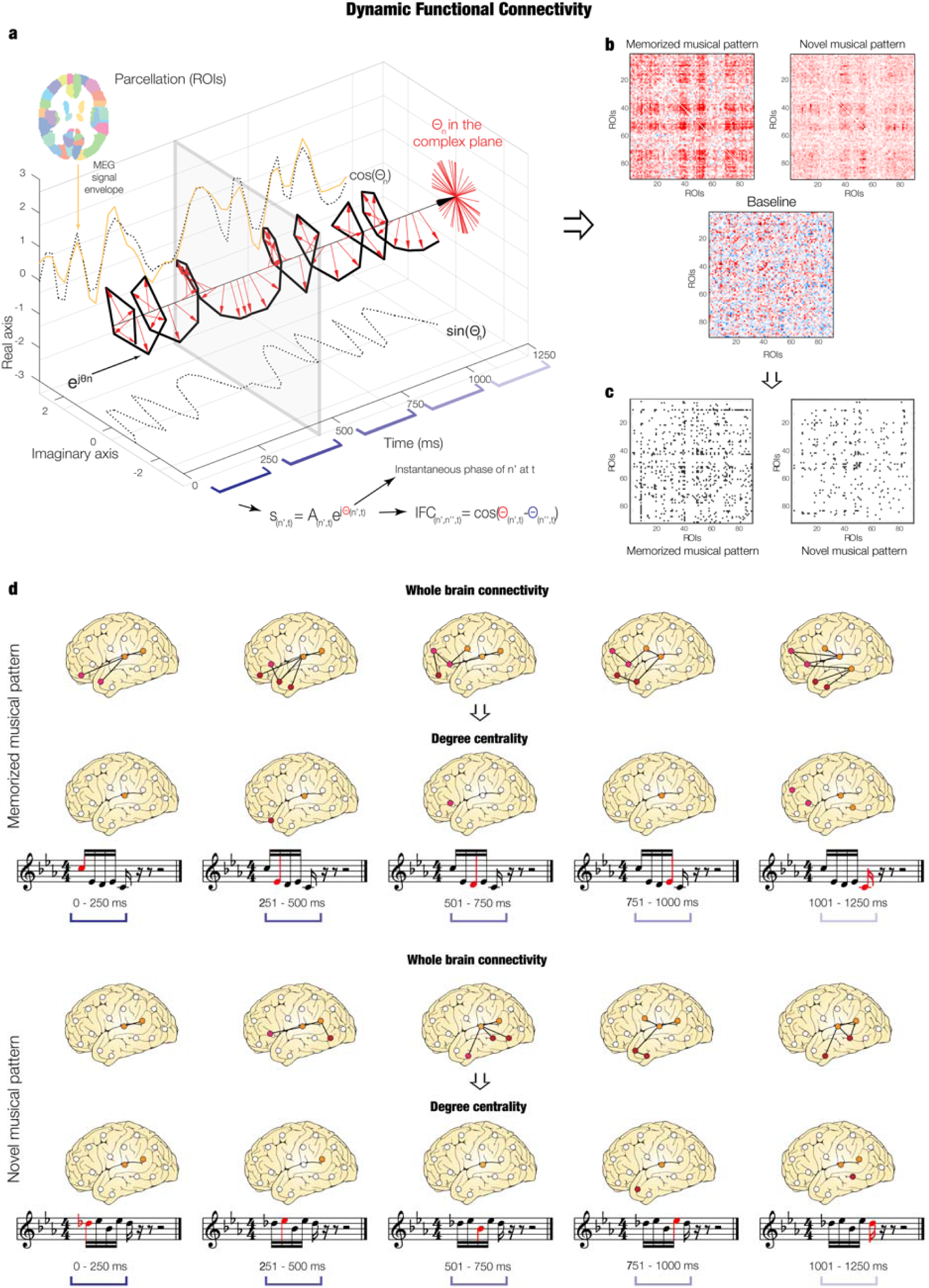
Overview of the analysis pipeline. **a -** We computed the Hilbert transform of the envelope of each AAL parcellation ROI and estimated the phase synchronization by calculating the cosine similarity between the instantaneous phases of each pair of ROIs. **b –** We obtained for each time-point two IFC matrices for the task: one for the memorized and one for the novel musical patterns, plus an additional one for resting state that was used as baseline in the following steps. **c –** We contrasted the task matrices versus the average of the baseline matrix to isolate the brain activity specifically related to the pattern recognition brain processes. **d –** We investigated the brain connectivity for each tone forming the musical sequences. First, we showed an illustration of the whole-brain connectivity dynamics. Then, we depicted the significantly central ROIs within the whole-brain network, estimated by applying MCS on the ROIs degree centrality calculated for each tone composing the musical sequences.

### MEG sensor data

Our first analysis was conducted on MEG sensor data and focused on the brain activity underlying the recognition of the previously memorized versus novel musical patterns. Specifically, we prepared 80 musical fragments comprising five notes with the same duration and lasting 1250 ms each (40 trials were excerpt of the Bach’s prelude, while the other 40 were novel melodies, matched in rhythm, volume, timbre, tempo, meter, tonality, information content and entropy, see Supplementary **Figure SF1** for all musical patterns). On average, participants correctly identified the 78.15 ± 13.56 % of the Bach’s excerpts (mean reaction times (RT): 1871 ± 209 ms) and the 81.43 ± 14.12 % of the novel melodies (mean RT: 1915 ± 135 ms). Subsequent MEG sensor data analysis was conducted on correct trials only.

#### Multivariate pattern analysis

As depicted in **Figure 3A**, we conducted a multivariate pattern analysis using a support vector machine (SVM) classifier (see details in the Methods section) to decode different neural activity associated with the recognition of previously memorized versus novel musical patterns. This analysis resulted in a decoding time series showing how neural activity differentiated the two experimental conditions. The decoding time series was significantly different from chance level in the time range 0.8 - 2.1 seconds from the onset of the first tone (*q* < .026, false-discovery rate (FDR)-corrected, **Figure 3A**, top left).

**Figure 3.**
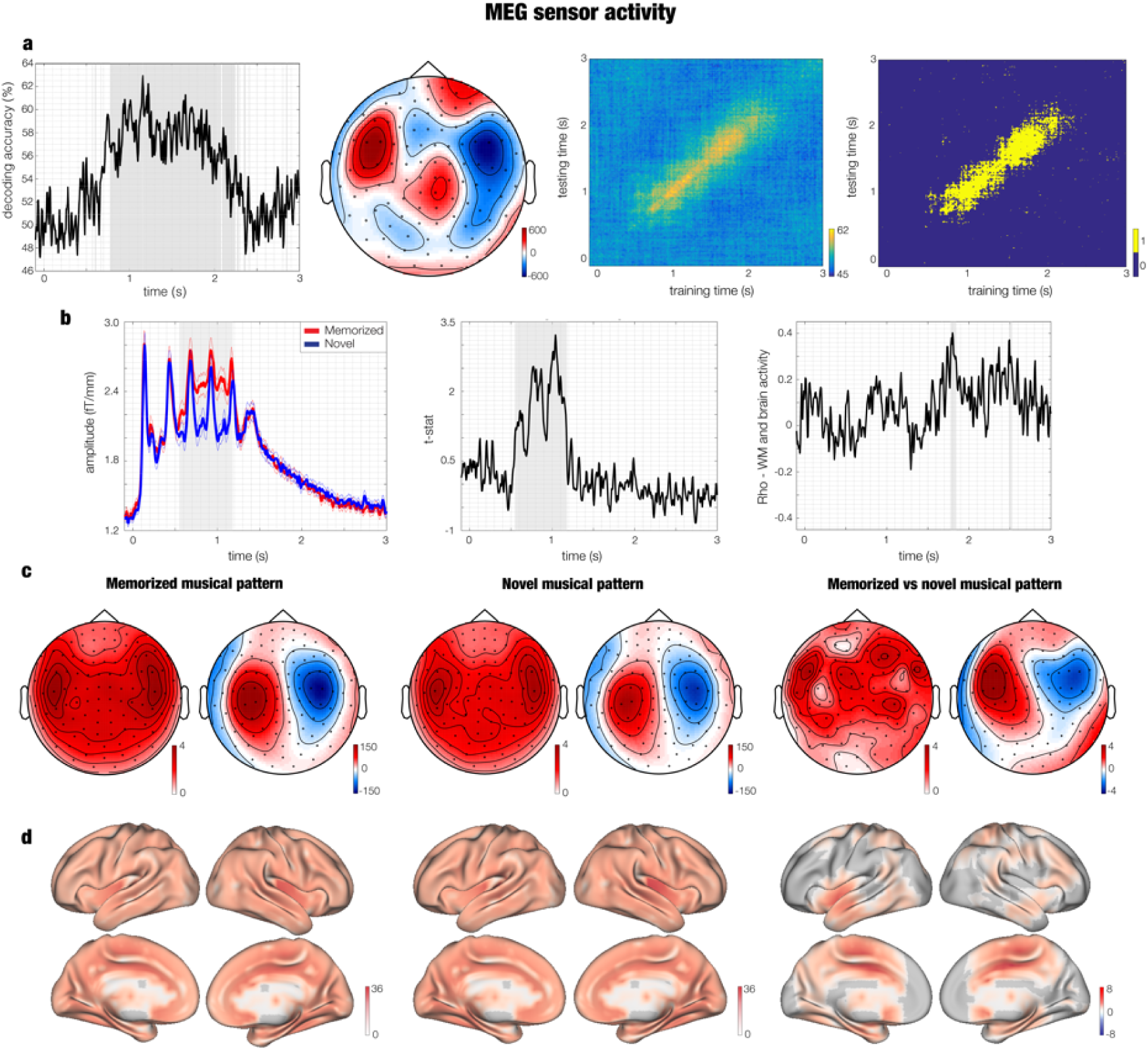
Brain activity underlying memorized versus novel musical patterns. **a –** Multivariate pattern analysis decoding the different neural activity associated to memorized versus novel musical patterns. Decoding time series (left), spatial patterns depicted as topoplot (middle left), temporal generalization decoding accuracy (middle right) and statistical output of significant prediction of training time on testing time (right). **b –** The left plot shows the amplitude associated to memorized (red) and novel musical patterns (blue). The middle plot illustrates the t-statistics related to the contrast between memorized versus novel musical patterns. The right plot shows the correlation between working memory (WM) abilities and the neural responses underlying recognition of the memorized versus novel musical patterns. Thinner lines depict standard errors. The plots refer to the average over the gradiometer channels forming the significant cluster outputted by MEG sensor MCS. **c –** Three couples of topoplots showing brain activity for gradiometers (left of each pair, fT/cm) and magnetometers (right of each pair, fT) within the significant time-window emerged from MCS. First couple of topoplots depicts the neural activity underlying the recognition of the previously memorized musical patterns, second couple refers to the novel musical patterns, while the third one represents the statistics (t-values) contrasting the brain activity underlying recognition of memorized versus novel musical patterns. **d –** Neural sources for the recognition of memorized patterns (left), novel patterns (middle) and their contrast (right). The values are t-statistics.

To evaluate the persistence of discriminable information over time, we applied a temporal generalization approach by training the SVM classifier at a given time point t, as before, but testing across all other time points. Intuitively, if representations are stable over time, the classifier should successfully discriminate signals not only at the trained time t, but also over extended periods of time that share the same neural representation. FDR-corrected (*q* < .005) results are depicted in **Figure 3A** (top right) showing that performance of the classifier was significantly above chance even a few hundreds of milliseconds beyond the diagonal. Nevertheless, our results showed that the strongest decoding values were on the main diagonal and the off-diagonal values decreased fast. As illustrated by previous research, this suggests that the difference between the two experimental conditions are likely to be mainly ballistic patterns (as shown by the close-to-the-diagonal decoding performance), perhaps leading to slightly more sustained and stable representations only at a later time, as shown in **Figure 3A** [19, 20].

#### Univariate tests and Monte Carlo simulations

The multivariate pattern analysis is a powerful tool that requires relatively few pre-processing steps for returning an estimation of neural activity that discriminates two or more experimental conditions. However, this technique does not provide directional information, that is, it does not identify which experimental condition yields a stronger neural signal. To answer this question, we performed independent univariate t-tests between conditions for each time-sample and each MEG channel and then corrected for multiple comparisons using a cluster-based Monte Carlo simulation (MCS) approach.

First, we contrasted the previously memorized versus novel musical patterns (t-test threshold = .01, MCS threshold = .001, 1000 permutations), considering the positive t-values only (which is when the memorized music was associated to a stronger brain activity than the novel melodies). We performed this analysis in the time-range 0 – 2.5 seconds by using combined planar gradiometers only. This procedure yielded the identification of one main significant cluster (MCS *p* < .001; time: 0.547 – 1.180 seconds, size: 2117), as depicted in **Figure 3B** and **3C** and reported in detail in **Tables ST1** and **ST3**. Then, on the basis of the significant cluster appearing, we computed the same algorithm one more time for magnetometers only, within the significant time-range emerged for the first MCS (0.547 – 1.180 seconds, *p* < .001, **Table ST1**). This two-step procedure was necessitated by the sign ambiguity typical of magnetometer data and returned three significant clusters (positive magnetometers: MCS *p* < .001; time: 0.627 – 1.180 seconds, size: 817: negative magnetometers: Cluster I - MCS *p* < .001; time: 0.727 – 0.880 seconds, size: 190; Cluster II - MCS *p* < .001; time: 0.960 – 1.133 seconds, size: 168).

Then, the same procedure was carried out by considering the results where the brain activity associated to the novel melodies exceeded the one elicited by Bach’s prelude excerpts. This analysis returned eight small significant clusters (size range: 6 – 14, *p* < .001) shown in **Table ST2**.

#### Relationship between brain activity and behavioral measures

Once we established that the recognition of the memorized and novel musical sequences gave rise to clearly different brain activity, we investigated whether such activity was modulated by individual differences such as working memory (WM) and musical skills related to the Bach’s prelude used in the study. Indeed, having better memory or musical skills, and a previous familiarity with the Bach’s piece could represent an advantage during the task and therefore should be considered. In our homogenous sample, only six of 70 participants declared to have previously played the Bach’s prelude used in the study in front of a public and importantly only four of them stated that they still remembered a few short bars. No one was sufficiently familiar with the prelude to be able to play it by heart. The rest of the participants reported low general familiarity with the piece in terms of listening habits (30 participants reported “*I have never heard the prelude before*”, 14 stated “*I have occasionally heard it”*, 12 declared “*I sometimes listen to it*”, two said “*I usually listen to it*”, three reported “*I played it for myself*”, six stated “*I played it in front of a public*”, three participants did not answer the question; see Methods for further details).

Using the averaged brain activity over the gradiometer channels forming the significant cluster identified with the univariate analysis, we computed the difference between the neural activity underlying recognition of memorized versus novel musical sequences. Then, we correlated each time-point of the resulting time series with four behavioral measures: WM skill (i), aesthetical judgement of the Bach’s prelude (ii), previous familiarity with the Bach’s prelude (iii), the Goldsmith Musical Sophistication Index (GOLD-MSI) (iv), which measures the ability of engaging with music. We corrected for multiple comparisons by using MCS with significance level α = .001. Overall, the results showed that the neural activity was not correlated to such measures. Indeed, detailed analysis revealed only few small significant clusters. Two clusters at the gradiometer level were found for WM (Cluster I: MCS *p* < .001; time: 1.77-1.85, mean r = 0.32; Cluster II: MCS *p* < .001; time: 2.48-2.53, mean r = 0.30, **Figure 3B**), meaning that participants with higher WM presented a stronger neural activity underlying musical recognition. Regarding liking, we detected two significant clusters (Cluster I: MCS p < .001; time: 2.40-2.45, mean r = 0.33 (**Figure SF2**); Cluster II: MCS p < .001; time: 1.11-1.14, mean r = -0.27). The two clusters were small and pointed to different directions of the correlation with the neural data (one was positive and the other negative), suggesting that the aesthetical judgment was not a good predictor of the brain activity. Finally, both familiarity with the Bach’s prelude and the Goldsmith musical index returned only one small cluster showing a negative correlation: Cluster I: MCS *p* < .001; time: 1.18-1.21, mean r = -0.28. (familiarity); Cluster I: MCS *p* < .001; time: 0.10-0.13, mean r = -0.29 (Goldsmith) (**Figure SF2**). No significant differences were detected when contrasting the brain activity underlying music recognition across pianists, non-pianist musicians and non-musicians (**Figure SF2**). Overall, these results show that aesthetic judgement, familiarity, and musical engagement have a negligible effect on the brain results.

### Source reconstructed data

To identify the neural sources of the signal, we employed a beamforming approach and computed a general linear model (GLM) for assessing, at each time-point, the independent neural activity associated to the two conditions as well as their contrasts.

#### Main cluster of previously memorized versus novel musical patterns

We identified the neural sources of the gradiometers significant cluster emerging from the MEG sensor data when contrasting memorized versus novel musical sequences. Here, we performed one permutation test in source space, with an α level of .05, which, in our case, corresponded to a cluster forming threshold of *t* = 1.7. As depicted in **Figure 3D**, results showed a strong activity originating in the primary auditory cortex, insula, hippocampus, frontal operculum, cingulate cortex and basal ganglia. Detailed statistics are provided in **Table ST4**.

#### Dynamic brain activity during development of musical patterns

To reveal the specific brain activity dynamics underlying the recognition of the musical sequences, we carried out a further analysis for each musical tone forming the musical pattern. Here, we adopted a stricter cluster forming threshold of *t* = 2.7 (see Methods for details). As depicted in **Figure 4A** and **4B**, we found significant activity within primary auditory cortex and insula, especially in the right hemisphere, for both experimental conditions. This activity decreased over time, following the unfolding of the musical sequences. Conversely, the contrast between memorized versus novel music gave rise to a burst of activity for the memorized Bach’s excerpts increasing over time, especially with regards to the last three tones of the musical sequences, as shown in **Figure 4C**. This activity was mainly localized within hippocampus, frontal operculum, cingulate cortex, insula, inferior temporal cortex and basal ganglia. We report detailed clusters statistics in **Table ST5**.

**Figure 4.**
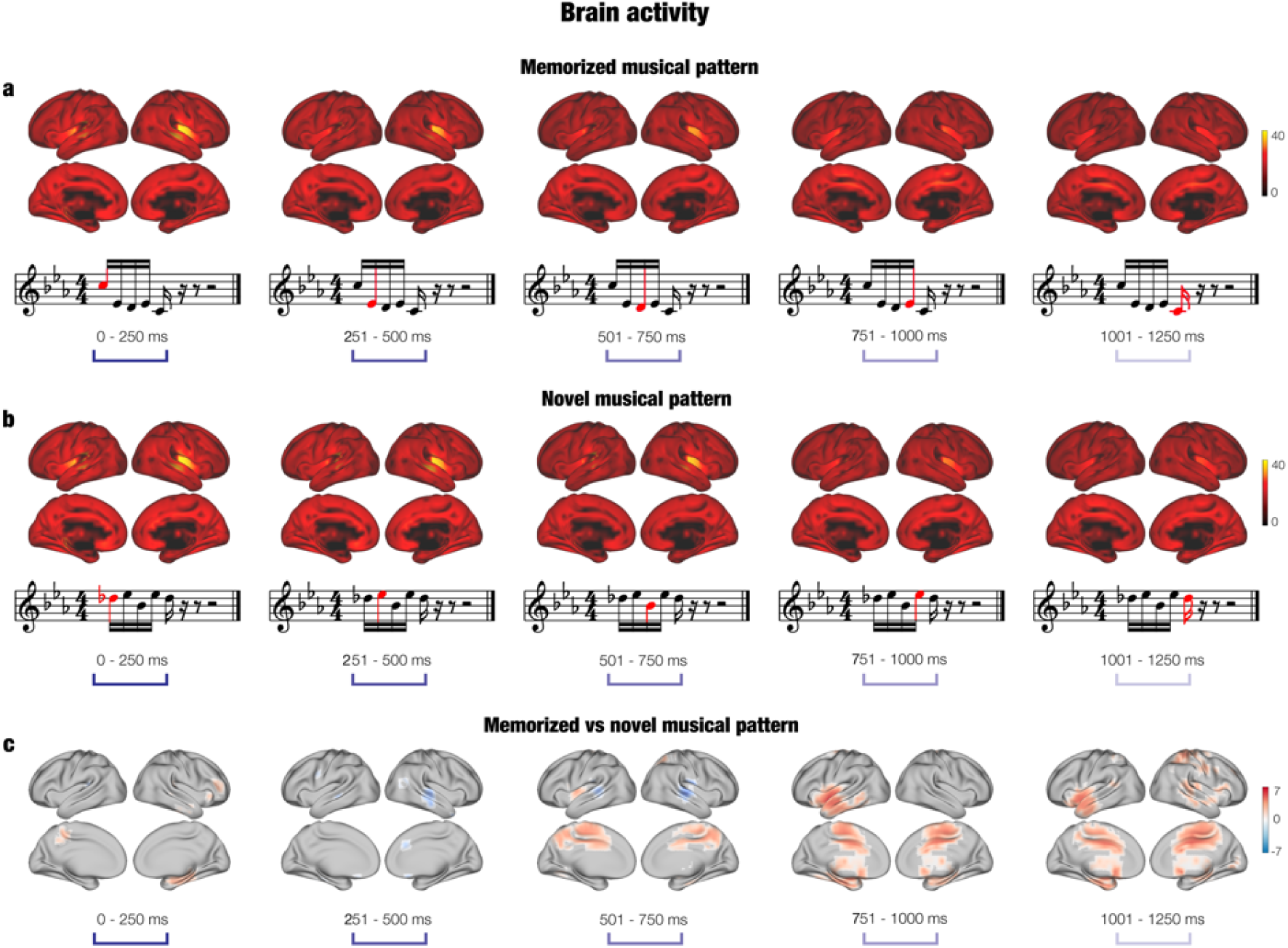
Brain activity over time. **a –** Brain activity (localized with beamforming) associated to the recognition of the previously memorized musical patterns (top row). Such patterns were extracted from the Bach’s prelude that participants attentively listened to before doing the recognition task. The bottom row depicts an example trial for the memorized patterns, extracted from Bach’s prelude). Red tones illustrate the dynamics of the musical excerpt. **b –** Brain activity underlying the detection of the novel musical pattern (top row) and musical representation of one example trial (bottom row). **c –** Contrast (t-values) over time between the brain activity underlying memorized versus novel musical patterns.

### Functional connectivity

To obtain a better understanding of the brain dynamics underlying recognition, we complemented our brain activity results with an investigation of the static and dynamic functional connectivity.

#### Static functional connectivity

MEG pre-processed data was constrained to the 90 non-cerebellar parcels of the automated anatomic labelling (AAL) parcellation and corrected for source leakage. First, we computed Pearson’s correlations between the envelopes of the time series of each pair of brain areas. This procedure was carried out for both task (in this case without distinguishing between memorized and novel music) and resting state (used as baseline) for each participant and five frequency bands: delta, theta, alpha, beta and gamma. Then, we tested the overall connectivity strengths of the five frequency bands during auditory recognition by employing analysis of variance (ANOVA). The test was significant (*F*(4,330) = 187.02, *p* < 1.0e-07). As depicted in **Figure 5A** and **5B**, post-hoc analysis highlighted especially that theta band had a stronger connectivity profile than all other frequency bands (*p* < 1.0e-07).

**Figure 5.**
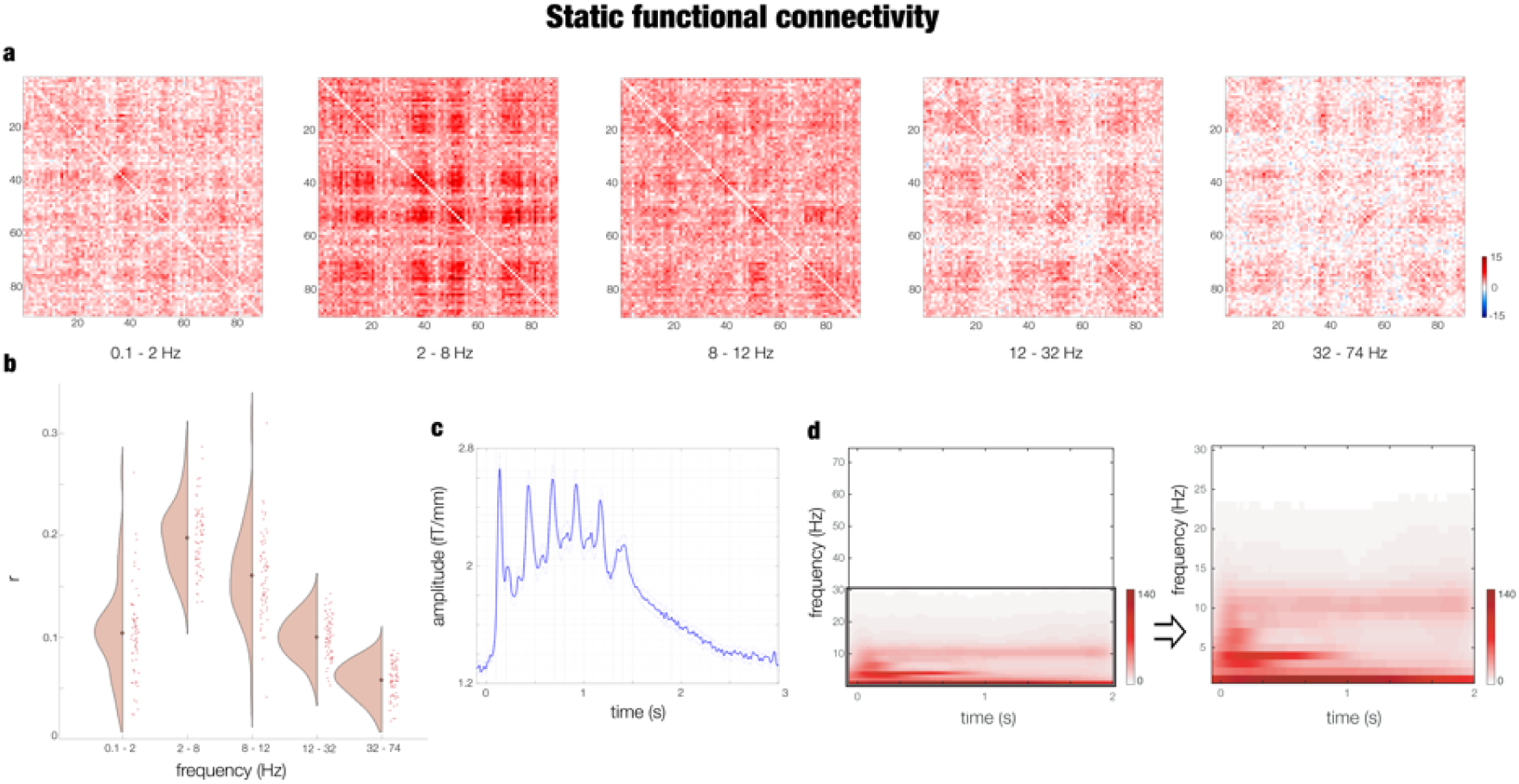
Static functional connectivity. **a –** Contrast between recognition task (memorized and novel musical patterns averaged together) and baseline SFC matrices calculated for five frequency bands: delta (0.1 – 2 Hz), theta (2 – 8 Hz), alpha (8 – 12 Hz), beta (12 – 32 Hz), gamma (32 – 74 Hz). **b –** Violin-scatter plot showing the average of the SFC matrices over their two dimensions for all participants. **c –** Averaged MEG gradiometer channels waveform of the brain activity associated to the recognition task. **d –** Power spectra for all MEG channels associated to the recognition task. The first power spectra matrix reflects the analysis from 1 to 74 Hz in 1-Hz intervals, while the second from 1 to 30 Hz in 1-Hz intervals.

To detect the significance of each brain region centrality within the whole-brain network for the auditory recognition task, we contrasted the brain connectivity matrices associated to the task versus baseline by performing a Wilcoxon signed-rank test for each pair of brain areas. Then, the resulting *z*-values matrix was submitted to a degree MCS (see Methods for details). We computed this analysis independently for the five frequency bands and therefore we considered significant the brain regions whose *p*-value was lower than the α level divided by 5 (2.0e-04). The results for theta band are depicted in **Figure SF3** and reported as follows: left Rolandic operculum (*p* < 1.0e-07), insula (*p* < 1.0e-07), hippocampus (*p* = 5.5e-05), putamen (*p* < 1.0e-07), pallidum (*p* < 1.0e-07), caudate (*p* = 1.1e-05), thalamus (*p* < 1.0e-07), Heschl’s gyrus (*p* < 1.0e-07), superior temporal gyrus (*p* < 1.0e-07), right superior temporal gyrus (*p* = 1.1e-06), Heschl’s gyrus (*p* < 1.0e-07), thalamus (*p* < 1.0e-07), parahippocampal gyrus (*p* = 4.3e-05), pallidum (*p* < 1.0e-07), putamen (*p* < 1.0e-07), amygdala (*p* < 1.0e-07), insula (*p* < 1.0e-07) and Rolandic operculum (*p* < 1.0e-07). Additional results related to the other frequency bands are reported in supporting information (SI) Appendix (**SR2**).

Conversely, the degree MCS of the contrasts between memorized versus novel melodies yielded no significant results.

#### Dynamic functional connectivity

Expanding on the analysis of static functional connectivity, we focused on the brain connectivity patterns evolving dynamically over time. Thus, by employing Hilbert transform and cosine similarity, we computed the phase synchronization between each pair of brain areas, obtaining one instantaneous functional connectivity (IFC) matrix for each musical tone (see Methods for details). Then, we calculated the contrast between the IFC matrices associated to the two conditions versus the baseline (illustrated in the top rows of **Figure 6A** and **6B**, and in **Figure SF4**). Subsequently, as reported in the middle rows of **Figure 6A** and **6B**, we computed the significantly central brain regions within the whole-brain network for each tone forming the musical pattern and both memorized and novel musical patterns. We tested statistical significance by using MCS. Since we repeated this test 10 times (two conditions and five tones), we used a threshold = 1.0e-04, obtained dividing the α level (.001) by 10. Detailed statistics are reported in **Table ST6**. Finally, as depicted in **Figure 6C** (top row), we contrasted the IFC matrices for memorized versus novel musical sequences and then we estimated the correspondent significantly central brain regions (**Figure 6C**, bottom row). These results showed a higher centrality for memory and evaluative regions such as cingulate, hippocampus and subgenual cortex for the memorized musical patterns, while the novel patterns were associated to an overall stronger centrality of auditory areas. Results are reported in detail in **Tables ST7** (memorized musical patterns)**, ST8** (novel musical patterns) and **SR3** (whole-brain phase synchronization coupling).

**Figure 6.**
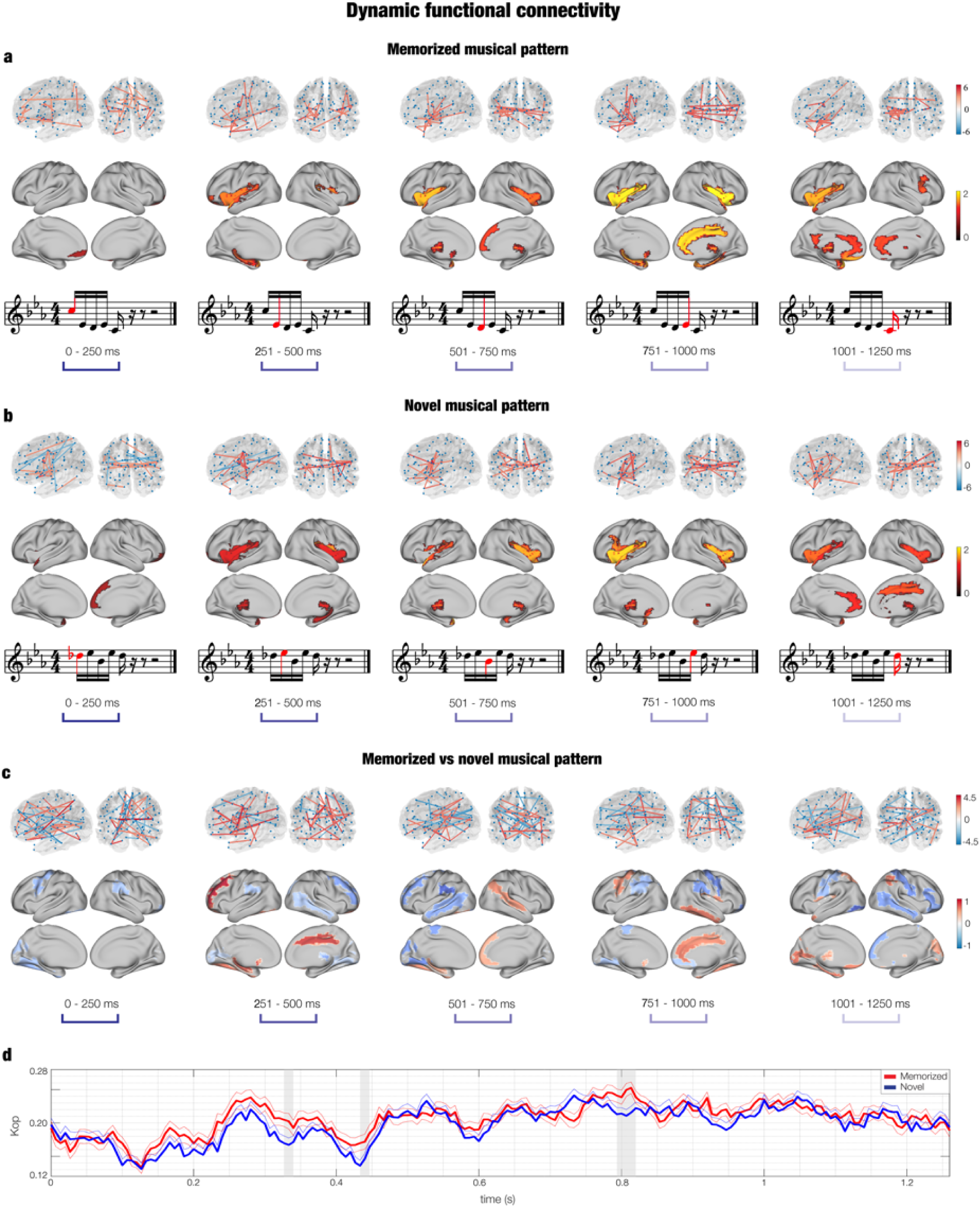
Dynamic functional connectivity. **a –** DFC calculated for each tone composing the memorized musical patterns. The top row depicts the whole-brain connectivity in a brain template (each pair presents the left hemisphere and a posterior view of the brain). Values refers to Wilcoxon sign-rank test z-values computed for the memorized musical patterns versus baseline DFC matrices. The bottom row illustrates the significantly central ROIs within the whole-brain network over time, as assessed by MCS. Values show the average z-value between the significant ROIs and the rest of the brain regions. **b –** DFC calculated for the novel musical patterns. The depiction is analogous to the one described for point a). **c –** DFC computed by contrasting memorized versus novel musical patterns. The depiction is analogous to the one described for point a). **d –** Kuramoto order parameter calculated for each time-point for both memorized (red line) and novel (blue line) musical patterns. Grey areas show the significant time-windows emerged by MCS.

### Kuramoto order parameter

Finally, to obtain a global measure of instantaneous connectivity within the brain over time, we computed the Kuramoto order parameter (see Methods) for the two experimental conditions and each time-point of the recognition task, obtaining two time series. We contrasted them by performing t-tests and corrected for multiple comparisons through a 1-dimensional (1D) MCS. Results, depicted in **Figure 6D**, showed a significant difference for the following time-ranges: 0.793 – 0.820 seconds (*p* = 4.5e-05), 0.327 – 0.340 seconds (*p* = .01) and 0.433 – 0.447 seconds (*p* = .01). In all three cases, the recognition of the memorized versus novel music was characterized by a significantly higher Kuramoto order parameter.

## Discussion

In this study, we were able to detect the fine-grained spatiotemporal dynamics of the brain activity and connectivity during recognition of previously memorized auditory patterns compared to carefully matched novel melodies.

First, by using a multivariate pattern analysis and MCS of massive univariate data, we found that the brain activity elicited by the recognition of excerpts from Bach’s prelude compared to the novel musical patterns gave rise to significant changes in widespread regions including primary auditory cortex, superior temporal gyrus, cingulate gyrus, hippocampus, basal ganglia, insula and frontal operculum. Notably, the neural difference reflecting the recognition of memorized versus novel musical patterns extended from approximately 700ms to 2000ms after the onset of the first tone of the melodies. This suggests that the brain discriminated the two categories of musical patterns, especially from tone number three of the sequences. Interestingly, this difference in neural activity extended up to 2000ms, which corresponded to about 100ms after participants categorized the musical patterns by using the response pad (the mean reaction time was approximately 1900ms for both categories of patterns). This evidence could indicate that the elaborated process of recognition and discrimination of the two categories of musical sequences requires a widespread network of brain areas whose activity was differentiated for more than 1000ms.

Second, we investigated this important finding further by estimating static and dynamic functional brain connectivity evolving over time. Here, the recognition of the musical patterns was accompanied by significant centrality within the whole-brain network of a number of brain regions including insula, hippocampus, cingulate gyrus, auditory cortex, basal ganglia, frontal operculum, subgenual and orbitofrontal cortices. Notably, these connectivity patterns emerged more clearly for the last three tones (out of the five tones) of the musical patterns and, even if similar across conditions, they were stronger for the memorized versus novel musical sequences. This finding lends strong support to the intriguing proposal by Changeux that recognition of meaningful temporal patterns, and more generally art, has the ability to ignite large brain networks comprising areas related to memory, evaluation, pleasure and abstract thinking [21]. As such, besides their primary significance in relation to memory, our results may also be seen with regards to affective neuroscience, suggesting that to allow recognition of previously memorized music, the brain also recruits areas usually associated to pleasure and eudaimonia [22].

The brain activity recorded during the recognition of the musical sequences was coherent with a large number of studies showing auditory processes associated to primary auditory cortex and insula [20, 21]. Remarkably, contrasting the previously memorized versus novel music, we observed stronger activity underlying the recognition of the memorized sequences in brain areas related to memory recognition such as hippocampus, medial temporal cortices [25, 26] and cingulate cortices [27]. Additionally, the recognition of excerpts from Bach’s prelude was associated to a stronger activity of brain regions previously related to evaluative processes [28, 29] and pleasure [30, 31] such as cingulate gyrus, subgenual cortices as well as parts of the basal ganglia. Finally, recognition of memorized music was accompanied by stronger activity in brain regions responsible for fine-grained auditory elaboration and prediction error such as the inferior temporal cortex [32] and insula [33, 24].

In addition to investigating the significant differences in brain activity ignited by recognition of previously memorized versus novel music, we conducted detailed functional connectivity analyses. Since music is a language that acquires meaning over time, we mainly focused on dynamic functional connectivity analyses to assess whether different connectivity patterns were associated to different parts of the musical sequence. Remarkably, our connectivity analysis revealed both important similarities and differences with the evolving brain activity over time.

A key relevant difference was that while the activity of the brain areas decreased over time, the degree centrality and connectivity between brain regions became clearer with the temporal development of the auditory patterns. Moreover, primary auditory cortex did not play a crucial role, while the main connectivity patterns emerged for brain regions related to higher sound and linguistic elaborations such as insula, inferior temporal cortex [33, 32] and frontal operculum [34]. Furthermore, we also observed other central brain areas that have been previously related to evaluative processes such as orbitofrontal and subgenual cortices [35, 28] and memory such as hippocampus [36] and basal ganglia [37]. Notably, the connectivity patterns and centrality of brain regions were stronger for the memorized compared to the novel musical patterns. This evidence suggests the relevance of the synchronization between brain areas over their mere activation to understand the brain networks underlying complex memory processes.

Conversely, a crucial similarity between brain activity and connectivity analyses was that compared to the novel melodies, the recognition of the memorized musical patterns ignited the shared common music processing brain network in terms of both activity and connectivity. As mentioned earlier, Changeux [21] proposed that processing and recognition of certain privileged classes of memorized stimuli, including meaningful temporal patterns and artistic works, can ignite the brain areas forming the global neuronal workspace [21]. Authors defined the global workspace as a privileged network of brain areas, where conscious information is processed in terms of memory, attention, and valence, and subsequently broadcast and made available to the whole-brain [38,39,40]. As predicted by Changeux’s hypothesis, the recognition of the memorized musical excerpts taken from Bach’s prelude – over and above the novel melodic patterns – led to stronger ignition of putative regions in the global workspace such as hippocampus, cingulate gyrus, orbitofrontal cortex, and frontal operculum, perhaps reflecting the mechanisms that allow the brain to process, extract a meaningful representation, and recognize previously memorized musical patterns. Interestingly, our findings showed that the conscious, effortful recognition of temporal patterns involved several high-order brain areas, while previous studies on automatic recognition and prediction error associated to sudden deviations in auditory sequences (e.g. indexed by MMN and N100) revealed a primary contribution of sensorial brain areas such as auditory cortices [10, 11]. This provides evidence for the relevance of the global neuronal workspace for conscious over automatic temporal pattern discrimination and recognition.

A further central theory in the neuroscientific field that can be related to our results is predictive coding. In this framework, the brain is considered a generator of models of expectations of the incoming stimuli. Recently, this theory has been linked to complex cognitive processes, finding a remarkable example in the neuroscience of music [2]. In their work, Koelsch and colleagues suggested that the perception of music is the result of an active listening process where individuals constantly formulate hypothesis about the upcoming development of musical sentences, while those sentences are actually evolving and unfolding their ambiguities. Our study is consistent with this perspective, where the brain predicts the upcoming sounds of the musical patterns leading to at least two separable outcomes. On the one hand, there is activity in the primary auditory cortex, responsible for the first sensorial processing of tones, and decreasing over time. On the other hand, the ignition of brain areas related to memory and evaluative processes is increasing over time and stronger for the recognition of the memorized versus novel musical sequences. Indeed, this would suggest that the brain has formulated predictions of the upcoming sounds based on the memory trace previously stored during the encoding part of our experimental task. The match between those predictions and the actual sounds presented to participants may lead to the ignition of the brain areas that we observed in our experiment.

In line with the predictive coding framework, our results also expanded the neuroscientific literature on prediction error. Indeed, previous research using paradigms that required active encoding and evaluation of the stimuli to formulate predictions and drive decision-making processes has highlighted a prominent role of the anterior cingulate gyrus [40, 41] and a relevant contribution of hippocampal areas [42, 43]. Furthermore, in the auditory domain, traditionally prediction error has been investigated through the brain response to violations of auditory regularity, as indexed by event-related components such as N100 and MMN [44, 10]. Additionally, more recent research has shown a hierarchical organization of prediction error processes occurring within the auditory cortex [45, 46]. However, the majority of these auditory studies employed varied versions of the oddball paradigm or more complex tasks that did not require conscious elaboration of the stimuli, making it difficult to interpret the results in terms of higher brain functioning.

In contrast, in the current study we utilized an auditory paradigm that required participants to make active choices based on prediction processes. Indeed, in line with the previous literature, we detected a prominent role played by cingulate cortex, hippocampus and auditory cortex to establish whether a musical excerpt was already known or was a novel sequence. Notably, we observed hippocampus and cingulate gyrus to be central within the whole-brain network when the incoming stimuli matched the prediction made by the brain (e.g. when listening to excerpts from Bach’s prelude), while the auditory cortex appeared more central and active for signalling a violation within the musical sequence expectation (e.g. when listening to novel melodies). Once again, our results suggest that while the automatic detection of regularities and deviations in temporal patterns relies on recruitment of modular, sensorial brain areas, the conscious, effortful recognition of meaningful temporal patterns recruits a widespread brain network largely overlapping with the previously described global neuronal workspace [21].

Additionally, to increase the reliability of our findings, we complemented our main results on spatiotemporal dynamics of brain activity and connectivity with more traditional approaches such as static functional connectivity and analysis of MEG sensors.

For the static functional connectivity, we computed Pearson’s correlations between the envelopes of each pair of brain areas and then studied the centrality of brain regions within whole-brain networks. We detected the most prominent connectivity patterns associated to music recognition for theta band (2 – 8 Hz), highlighting a large network of brain regions including primary auditory cortex, superior temporal gyrus, frontal operculum, insula, hippocampus, and basal ganglia. As expected, the static functional connectivity network was very similar to the dynamic functional connectivity ones and involved brain regions previously related to auditory [47], memory [25, 26] and evaluative processes [29].

Importantly, the static functional connectivity analysis failed to detect differences between the previously memorized and novel musical patterns. This shows that to capture the fine-grained brain functioning underlying recognition of memorized musical patterns, it was crucial to employ methods designed to detect the dynamic unfolding of the brain connectivity.

We also investigated our data by focusing on MEG sensor analysis. In this regard, we found that brain activity was reflected by two event-related field (ERF) components: N100 to each sound and a slow negativity following the entire duration of the musical patterns.

While the N100 associated to each tone represented a well-known finding, to the best of our knowledge the detection of the slow negative component associated to pattern recognition has never been described earlier. Indeed, even if this negative waveform shared similarities with well-established ERF components such as contingent negative variation (CNV) [48], P300 and P600 [49], previous literature has never associated an auditory recognition task to a slow negative component such as the one that we described. Thus, our results provided new insights and may contribute to develop future perspectives also within the framework of ERF and MEG sensor analysis.

Finally, analyzing the relationship between the brain activity during our recognition task and behavioral measures related to memory and musical skills showed very weak associations in small, isolated clusters. This suggested that having more engagement with music, general musical expertise, or a previous familiarity and higher appreciation for the Bach’s prelude does not play a major role in modulating the brain activity during the musical recognition task. Notably, in general our sample comprised homogeneous participants who did not have a wide previous familiarity nor knowledge of the Bach’s prelude. This does not, of course, rule out the hypothesis that prior knowledge of the music could modulate the brain activity during its recognition. However, our results strongly suggest that a previous vague familiarity with the piece is not enough to modulate brain functioning during musical recognition. Notably, a mild yet interesting effect was observed for WM. This evidence, coherently with previous research [50], shows a connection between WM skills and neural data underlying memory tasks, indicating that the brain of individuals with higher WM abilities is characterized by a stronger activity when recognizing temporal patterns such as the excerpts from Bach’s prelude. Our results suggest that memory skills may be more important than musical abilities and expertise when recognizing temporal sequences, even when they consist of musical melodies.

In conclusion, our findings have identified the spatiotemporal unfolding of fast-scale brain activity and functional connectivity elicited by the recognition of previously memorized compared to novel musical patterns. Since music is an excellent example of a language which gains meaning through the combination of its constituent elements over time, this study shed new light on the brain mechanisms underlying the processing and understanding of temporal patterns. Furthermore, by using state-of-the-art neuroimaging and analysis methods, our results highlight how the integration of brain activity and fast-scale phase synchronization analyses, combined with hierarchically organized musical patterns, provide a unique opportunity to understand the brain functioning underlying complex, cognitive processes such as temporal patterns recognition.

## Methods

### Data and code availability

The dataset and codes generated during this study are available on GitHub (https://github.com/leonardob92/LBPD-1.0.git). Please, contact the author Leonardo Bonetti for further information.

### Participants

The study comprised 70 volunteers coming from different Western countries and living in Denmark at the time of the experiment. Thirty-six of them were males and 34 were females (age range: 18 – 42 years old, mean age: 25.06 ± 4.11 years). Since our experiment involved a musical piece usually played by classical pianists, we recruited 23 classical pianists (13 males and 10 females, age range: 18 – 34 years old, mean age: 24.83 ± 4.10 years old), 24 non-pianist musicians (12 males and 12 females, age range: 19 - 42 years old, mean age: 24.54 ± 4.75), and 23 non-musicians (11 males and 12 females, age range: 21 – 35 years old; mean age: 25.86 ± 3.34). The sample regarding functional connectivity analysis slightly differed (three participants had to be discarded due to technical problems during acquisition) consisting of 67 participants (34 males and 33 females, age range: 18 – 42 years old, mean age: 25.00 ± 4.18 years). Specifically, 21 were non-pianist musicians (10 males and 11 females, age range: 42 – 19 years old, mean age: 24.29 ± 5.02 years), 23 classical pianists (13 males and 10 females, age range: 18 – 34 years old, mean age: 24.83 ± 4.10 years) and 23 non-musicians (11 males and 12 females, age range: 21 – 35 years old; mean age: 25.86 ± 3.34 years). Participants had homogeneous socio-economic and educational backgrounds and signed the informed consent before the beginning of the experiment.

All the experimental procedures complied with the Declaration of Helsinki – Ethical Principles for Medical Research and were approved by the Ethics Committee of the Central Denmark Region (De Videnskabsetiske Komitéer for Region Midtjylland) (Ref 1-10-72-411-17).

### Experimental design and stimuli

To study the brain dynamics of musical pattern recognition, we employed an old/new [51] auditory pattern recognition task during MEG recording (**Figure 1A)**. First, participants were requested to listen to four repetitions of a MIDI version of the right-hand part of the entire prelude in C minor BWV 847 composed by J.S. Bach. The tones had the same duration, which was of approximately 250ms. The full piece lasted about 2.5 minutes; thus, the total duration of the learning part was approximately 10 minutes (2.5 minutes repeated four times). Participants were asked to focus on the musical prelude and memorize it as much as possible. Second, they were presented with 80 short musical excerpts lasting 1250 ms each and requested to indicate whether each excerpt belonged to the prelude by Bach (memorized musical pattern, ‘old’, 40 trials) or was a novel musical pattern (‘new’, 40 trials). Subsequent analyses were performed on correctly recognized trials only. Importantly, the two categories of stimuli (memorized and novel musical patterns) were composed to be clearly distinguishable in the recognition task, even if they were matched among several variables, to prevent for potential confounds. Specifically, the two categories were matched for rhythm, volume, timbre, tempo, meter, tonality, information content (*IC*) and entropy (*H*). The memorized melodies consisted of excerpts of the Bach’s prelude. Specifically, we extracted one excerpt per musical bar, corresponding to the first five notes of the bar. These different excerpts were chosen since they were representative of the melodic contour and of the general repetitive structure of the Bach’s prelude. The novel musical sequences were created by using a melodic contour and intervals between the notes of the melodies that were different from Bach’s prelude excerpts. By doing so, we designed a task that was challenging yet feasible, since the two categories of melodies presented several similarities, yet were clearly different, since the melodic contours of the sequences and intervals between the tones were diverse in the two conditions. The 80 musical patterns are reported in musical notation in **Figure SF1**.

The *IC* and *H* were estimated for each tone of the Bach’s excerpts (mean *IC* : 5.70 ± 1.73, mean *H* : 4.70 ± .33) and of the novel melodies (mean *IC* : 5.92 ± 1.81, mean *H* : 4.78 ± .35) by using Information Dynamics of Music (IDyOM) [52]. This robust method uses machine learning to return a value of for the target note on the basis of a combination of the preceding notes of the musical piece comprising the target note and of a set of rules learned from a large set of prototypical pieces of Western music. Thus, in our study, the of each note of the musical patterns was computed using a model trained on both the Bach’s prelude excerpts and the novel melodies *(i)* and on the large corpus of prototypical pieces of Western music usually employed by IDyOM [52] *(ii)*. In this way, musical patterns of the two categories (memorized and novel patterns) with the same *IC* were composed of a series of intervals and melodic contours that were quite similar, and equally plausible in light of prototypical Western music.

Formally, the *IC* represents the minimum number of bits required to encode *e_i_* and is described by the equation (1):

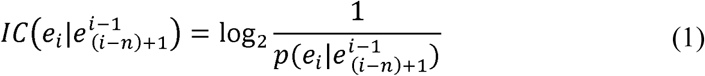

Where 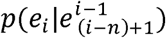 is the probability of the event *e_i_* given a previous set of 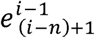 events.

The entropy gives a measure of the certainty/uncertainty of the upcoming event given the previous set of 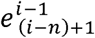 events and is calculated by the equation (2):

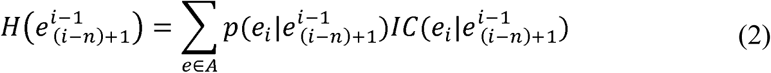

Equation (2) shows that if the probability of a given event *e_i_* is 1, the probability of the other events in *A* will be 0 and therefore *H* will be equal to 0 (maximum certainty). On the contrary, if all the events are equally likely, *H* will be maximum (maximum uncertainty). Therefore, IDyOM returns an estimation of the predictability of each tone and uncertainty with which it can be predicted, coherently with the human perception [53].

The entire prelude and the musical excerpts were created by using Finale (MakeMusic, Boulder, CO) and then presented by adopting Presentation software (Neurobehavioural Systems, Berkeley, CA).

We collected structural images for each participant by employing magnetic resonance imaging (MRI), either on the same day as the functional MEG scan or on another day within one month.

### Data acquisition

We acquired both MRI and MEG data in two independent sessions. The MEG data was acquired by employing an Elekta Neuromag TRIUX system (Elekta Neuromag, Helsinki, Finland) equipped with 306 channels. The machine was positioned in a magnetically shielded room at Aarhus University Hospital, Denmark. Data was recorded at a sampling rate of 1000 Hz with an analogue filtering of 0.1–330 Hz. Prior to the measurements, we accommodated the sound volume at 50 dB above the minimum hearing threshold of each participant. Moreover, by utilizing a 3D digitizer (Polhemus Fastrak, Colchester, VT, USA), we registered the participant’s head shape and the position of four headcoils, with respect to three anatomical landmarks (nasion, and left and right preauricular locations). This information was subsequently used to co-register the MEG data with the anatomical structure collected by the MRI scanner. The location of the headcoils was registered during the entire recording by using a continuous head position identification (cHPI), allowing us to track the exact head location within the MEG scanner at each time-point. We utilized this data to perform an accurate movement correction at a later stage of data analysis.

The recorded MRI data corresponded to structural T1. The acquisition parameters for the scan were: voxel size = 1.0 x 1.0 x 1.0 mm (or 1.0 mm^3^); reconstructed matrix size 256×256; echo time (TE) of 2.96 ms and repetition time (TR) of 5000 ms and a bandwidth of 240 Hz/Px. Each individual T1-weighted MRI scan was subsequently co-registered to the standard Montreal Neurological Institute (MNI) brain template through an affine transformation and then referenced to the MEG sensors space by using the Polhemus head shape data and the three fiducial points measured during the MEG session.

### Data pre-processing

The raw MEG sensor data (204 planar gradiometers and 102 magnetometers) was pre-processed by MaxFilter [53] for attenuating the interference originated outside the scalp by applying signal space separation. Within the same session, Maxfilter also adjusted the signal for head movement and down-sampled it from 1000 Hz to 250 Hz.

The data was converted into the Statistical Parametric Mapping (SPM) format and further analyzed in Matlab (MathWorks, Natick, Massachusetts, United States of America) by using Oxford Centre for Human Brain Activity Software Library (OSL) [54], a freely available toolbox that combines in-house-built functions with existing tools from the FMRIB Software Library (FSL) [55], SPM [56] and Fieldtrip [57]. The data was then high-pass filtered (0.1 Hz threshold) to remove frequencies that were too low for being originated by the brain. We also applied a 48-52 Hz notch filter to correct for possible interference of the electric current. The data was further downsampled to 150 Hz and few segments of the data, contaminated by large artefacts, were removed after visual inspection. Then, to discard the interference of eyeblinks and heart-beat artefacts from the brain data, we performed independent component analysis (ICA) to decompose the original signal in independent components. Then, we isolated and discarded the components that picked up eyeblink and heart-beat activities, rebuilding the signal from the remaining components [58]. The data was epoched in 80 trials (one for each musical excerpt) lasting 3500 ms each (100ms of pre-stimulus time), and low-pass filtered (40Hz threshold) to improve the subsequent analysis.

Then, correctly identified trials were analyzed by employing two different methodologies (multivariate pattern analysis and cluster-based MCS of independent univariate analyses) to strengthen the reliability of the results as well as broaden the amount of information derived by the data.

### Multivariate pattern analysis

We conducted a multivariate pattern analysis to decode different neural activity associated to the recognition of previously memorized versus novel musical patterns. Specifically, we employed SVMs [59], analyzing each participant independently. MEG data was arranged in a 3D matrix (channels x time-points x trials) and submitted to the supervised learning algorithm. To avoid overfitting, we employed a leave-one-out cross-validation approach to train the SVM classifier to decode the two conditions. This procedure consisted of dividing the trials into *N* different groups (here *N* = 8) and, for each time point, assigning *N – 1* groups to the training set and the remaining *N^th^* group to the testing set. Then, the performance of the classifier to separate the two conditions was evaluated. This process was carried out 100 times with random reassignment of the data to training and testing sets. Finally, the decoding accuracy time series were averaged together to obtain a final time series reflecting the performance of the classifier for each participant.

Then, to identify the channels that were carrying the highest amount of information required for decoding the two experimental conditions, we followed the procedure described by Haufe and colleagues [60] and computed the decoding patterns from the weights returned by the SVM.

Finally, to assess whether the two experimental conditions were differentiated by neural patterns stable over time, we performed a temporal generalization multivariate analysis. The algorithm was the same as the one described above, with the difference that in this case we used each time-point of the training set to predict not only the same time-point in the testing set, but all time-points [61,19,20].

In both cases, to test whether the decoding results were significantly different from the chance level (50%), we used a sign permutation tests against the chance level for each time-point and then corrected for multiple comparisons by applying FDR correction (α = .05; FDR-adjusted *q* < .026 for non-temporal generalization results and α = .02; FDR-adjusted *q* < .005 for temporal generalization results).

### Univariate tests and Monte Carlo simulations

The multivariate pattern analysis is a powerful tool that requires relatively few pre-processing steps for returning an estimation of the different neural activity associated to two or more experimental conditions. However, this technique does not identify which condition was stronger than the other nor the polarity of the neural signal characterizing the experimental conditions. To answer these questions and strengthen our results, we employed a different approach by calculating several univariate t-tests and then correcting for multiple comparisons by using MCS.

Before computing the t-tests, in accordance with a large number of other MEG and electroencephalography (EEG) task studies [62], we averaged the trials over conditions, obtaining two mean trials, one for the memorized and one for the novel musical patterns. Then, we combined each pair of planar gradiometers by sum-root square. Afterwards, we computed a t-test for each MEG channel and each time-point in the time-range 0 – 2.500 seconds, contrasting the two experimental conditions. Independently for the two MEG sensor categories, we reshaped the matrix for obtaining, for each time-point, a 2D approximation of the MEG channels layout that we binarized according to the *p*-values obtained from the previous t-tests (threshold = .01) and the sign of *t*-values. The resulting 3D matrix (*M*) was therefore composed by 0s when the t-test was not significant and 1s when it was. Then, to correct for multiple comparisons, we identified the clusters of 1s and assessed their significance by running MCS. Specifically, we made 1000 permutations of the elements of the original binary matrix *M*, identified the maximum cluster size of 1s and built the distribution of the 1000 maximum cluster sizes. Finally, we considered significant the original clusters that had a size bigger than the 99.9% maximum cluster sizes of the permuted data. The whole MCS procedure was performed for gradiometers and magnetometers (in the significant time-window emerged from gradiometers, see SI Appendix **SR1** for details), for memorized versus novel musical patterns and vice versa.

### Relationship between neural activity and behavioral measures

We investigated whether the neural activity underlying the musical recognition task was modulated by individual differences along the following four behavioral measures: WM skills (i), aesthetical judgement of Bach’s prelude used in the study (ii), previous familiarity with Bach’s prelude (iii), the Goldsmith Musical Sophistication Index (GOLD-MSI) (iv) [63], which measures the ability of engaging with music. Indeed, possessing better memory or musical skills and having a previous familiarity or knowledge of the Bach’s musical piece could have represented an advantage during the task and therefore we took it into account in our analyses. Regarding WM, we used the widely adopted Wechsler Adult Intelligence Scale (WAIS-IV) [64], which returned a reliable measure of individual WM. With regards to aesthetical judgement of Bach’s prelude, we utilized a 7-score Likert scale from -3 (very unpleasant) to +3 (very pleasant). Previous familiarity with Bach’s prelude was assessed asking participants to mark the number corresponding to one of the following options: *1) I have never heard it before; 2) I have occasionally heard it; 3) I sometimes listen to it; 4) I usually listen to it; 5) I played it for myself; 6) I played it in front of a public*. Further, in our sample, only six participants declared to have previously played the Bach’s prelude that we used in the study in front of a public and sole four of them stated that they still remembered a few bars of it.

Using the averaged brain activity over the gradiometer channels forming the significant cluster emerged from the univariate analyses described in the previous paragraph, we computed the difference between the neural activity underlying recognition of memorized versus novel musical patterns. Then, we computed Pearson’s correlations for each time-point of the resulting time series and the four behavioral measures described above. The time series obtained from the correlations were binarized according to the outcome of the tests (one was assigned to the significant correlations (*p* < .05), while zero to the non-significant ones). Those values were submitted to a one-dimensional MCS with an α level = .001, to correct for multiple comparisons. First, we extracted the clusters of the significant results emerged from the correlations (here, cluster means group of contiguous significant values (ones)). Then, we computed 1000 permutations and for each of them, we randomized the order of the binarized values and computed the maximum cluster size of the detected clusters of significant values. Finally, we built a reference distribution of the 1000 maximum cluster sizes (obtained from the 1000 permutations) and considered significant the original clusters that were larger than the 99.9% of the permuted ones.

### Source reconstruction

#### Beamforming

The brain activity collected on the scalp by MEG channels was reconstructed in source space using OSL [54] to apply a local-spheres forward model and a beamformer approach as inverse method [65] (**Figure 1B** and **1C**). We used an 8-mm grid and both magnetometers and planar gradiometers. The spheres model depicted the MNI-co-registered anatomy as a simplified geometric model, fitting a sphere separately for each sensor [65]. The beamforming utilized a diverse set of weights sequentially applied to the source locations for isolating the contribution of each source to the activity recorded by the MEG channels for each time-point [65, 66].

#### General Linear Model

An independent GLM was calculated sequentially for each time-point at each dipole location, and for each experimental condition [67]. At first, reconstructed data was tested against its own baseline to calculate the statistics of neural sources of the two conditions memorized and novel musical patterns. Then, after computing the absolute value of the reconstructed time series to avoid sign ambiguity of the neural signal, first-level analysis was conducted, calculating contrast of parameter estimates (memorized versus novel musical patterns) for each dipole and each time-point. Those results were submitted to a second-level analysis, using one-sample t-tests with spatially smoothed variance obtained with a Gaussian kernel (full-width at half-maximum: 50 mm).

Then, to correct for multiple comparisons, a cluster-based permutation test [67] with 5000 permutations was computed on second-level analysis results, taking into account the significant time-range emerged from the MEG sensors MCS significant gradiometer cluster. Therefore, we performed one permutation test on source space, using an α level of .05, corresponding to a cluster forming threshold of *t* = 1.7.

#### Brain activity underlying musical patterns development

Then, as depicted in **Figure 1D**, we performed an additional analysis considering the brain activity underlying the processing of each tone forming the musical sequences. To do that we computed a GLM for each time-point and source location and then corrected for multiple comparisons with a cluster-based permutation test, as described above [67]. Here, when computing the significant clusters of brain activation independently for the two experimental conditions (memorized and novel musical patterns), we computed 10 permutation tests on source space, adjusting the α level to .005 (.05/10), corresponding to a cluster forming threshold of *t* = 2.7. Regarding memorized versus novel musical patterns, we performed five tests and therefore the α level became .01 (.05/5), corresponding to a cluster forming threshold of *t* = 2.3.

### Functional connectivity pre-processing

Regarding functional connectivity analysis, we used slightly larger epochs (pre-stimulus time of 200ms instead of 100ms). After reconstructing the data into source space, we constrained the beamforming results into the 90 non-cerebellar regions of the AAL parcellation, a widely-used and freely available template [68] in line with previous MEG studies [69] and corrected for source leakage [70]. Finally, according to a large number of MEG and EEG task studies [58] we averaged the trials over conditions, obtaining two mean trials, one for memorized and one for novel musical patterns. In order to minimize the probability of analyzing trials that were correctly recognized by chance, here we only considered the 20 fastest (mean RT: 1770 ± 352 ms) correctly recognized previously memorized musical patterns (mean RT: 1717 ± 381 ms) and novel musical patterns (mean RT: 1822 ± 323 ms) excerpts. The same operation has been carried out for the resting state that served as baseline. Here, we created 80 pseudo-trials with the same length of the real ones, starting at random time-points of the recorded resting state data.

This procedure has been carried out for five different frequency bands (delta: 0.1 – 2 Hz, theta: 2 – 8 Hz alpha: 8 – 12 Hz, beta: 12 – 32 Hz, gamma: 32 – 75 Hz) [71].

### Static functional connectivity and degree centrality

We estimated the SFC calculating Pearson’s correlations between the envelope of each pair of brain areas time courses. This procedure has been carried out for both task and baseline and for each of the five frequency bands considered in the study. Afterwards, we averaged the connectivity matrices in order to obtain one global value of connectivity for each participant and each frequency band. These values were submitted to an ANOVA to highlight which frequency band yielded the strongest connectivity effects, as illustrated in **Figure 5B** [72]. A follow up post-hoc analysis was conducted using Tukey’s correction for multiple comparisons. Then, for all frequency bands, we computed Wilcoxon sign-rank tests comparing each pair of brain areas for recognition task versus baseline, aiming to identify the functional connectivity specifically associated to the task. To assess the resulting connectivity matrix *B*, we identified the degree of each region of interest (ROI) and tested its significance through MCS [73].

In graph theory the weighted degree of each vertex *v* (here each brain area) of the graph *G* (here the matrix *B*) is given by the sum of the connection strengths of *v* with the other vertexes of *G*, showing the centrality of each *v* in *G* [74]. We computed the degree of each vertex of *B* for each musical tone, obtaining a 90 x 1 vector (*s_t_*). Then, through MCS, we assessed whether the vertices of *B* had a significantly higher degree than the degrees obtained permuting the elements of *B*. Specifically, we made 1000 permutations of the elements in the upper triangle of *B* and we calculated a 90 x 1 vector *d_v,p_* containing the degree of each vertex *v* for each permutation *p*. Combining vectors *d_v,p_* we obtained the distribution of the degrees calculated for each permutation. Finally, we considered significant the original degrees stored in *s_t_* that randomly occurred during the 1000 permutations less than two times. This threshold was obtained by dividing the α level (.001) by the five frequency bands considered in the study. The α level was set to .001 since this is the threshold that, during simulations with input matrices of uniformly distributed random numbers, provided no false positives. This procedure was carried out for each frequency band and for both experimental conditions.

### Phase synchronization estimation

In order to reveal the brain dynamics of the musical sequence recognition, we studied the phase synchronization between brain areas for theta, the frequency band that showed the strongest connectivity patterns when contrasting task versus baseline at the previous stage of analysis.

As illustrated in **Figure 2A**, by applying the Hilbert transform on the envelope of the reconstructed time-courses we obtained the analytic signal expressed by the following equation:

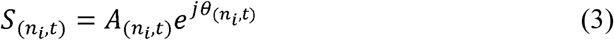

where *A_(ni,t)_* refers to the instantaneous amplitude and *θ_(ni,t)_* to the instantaneous phase of the signal for the brain region *n_i_* at time *t*. To prevent boundary artefacts due to the instantaneous phase estimation [75], we computed the Hilbert transform on time series that were slightly larger than the duration of the stimuli and then discarded their edges. To estimate the phase synchronization between two brain areas *n_i_* and *n_m_*, after extracting the instantaneous phase at time *t*, we calculated the cosine similarity expressed by equation (4):

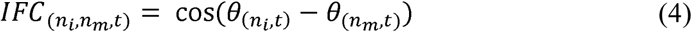

We carried out this procedure for each time-point and each pair of brain areas, obtaining 90 x 90 symmetric IFC matrices showing the phase synchronization of every couple of brain areas over time. This calculation has been performed for the two conditions (memorized and novel musical patterns) and for the baseline, as depicted in **Figure 2B**.

### Dynamic functional connectivity and degree centrality

Since we were interested in detecting the brain connectivity profile and dynamics associated to the processing of each tone of the musical sequence, we sub-averaged the IFC matrices to obtain one sub-averaged IFC matrix for each tone and condition. This procedure allowed us to reduce data dimensionality and suppress ongoing random noise.

To detect the brain connectivity specifically associated to the task we applied Wilcoxon signed-rank tests, contrasting, for each pair of brain areas, each of the five sub-averaged IFC matrices of the task with the averaged values over time of the baseline, as depicted in **Figure 2C**. This procedure yielded a square matrix for each tone (*B_t_*) and for both conditions (memorized and novel musical patterns) showing the strength of the phase synchronization between each pair of brain areas during the evolution of the task. After this procedure, we calculated an MCS (**Figure SF6**) analogous to the one described above for the SFC to test which ROIs were significantly central within the brain network for the recognition of memorized and novel musical patterns. Since we repeated this test 10 times (two conditions and five tones), we used a threshold = 1.0e-04, obtained by dividing the α level (.001) by 10. **Figure 2D** provides a graphical representation of the whole-brain connectivity and the correspondent degree centrality calculation. Finally, we contrasted the IFC matrices of memorized versus novel musical patterns and vice versa and we submitted the results to MCS. In this case, since we computed five independent tests (memorized versus novel musical patterns and novel versus memorized musical patterns), we obtained a threshold = 2.0e-04 (α level divided by five). We indicated the output of this calculation instantaneous brain degree (*IBD_(t)_*).

### Kuramoto order parameter

In conclusion, we computed the Kuramoto order parameter to estimate the global synchronization between brain areas over time. This parameter is defined by equation (5):

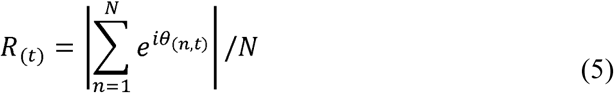

where *θ_(n,t)_* is the instantaneous phase of the signal for the brain region *n* at time *t* . The Kuramoto order parameter indicates the global level of synchronization of a group of *N* oscillating signals [76]. Thus, if the signals are completely independent the *N* instantaneous phases are uniformly distributed and therefore *R_(t)_* tends to 0. In contrast, if the *N* phases are equal, *R_(t)_* = 1.

The time series obtained for the recognition of memorized and novel musical patterns were then contrasted by calculating t-tests and binarized, assigning 1 to significant values (threshold = .05) emerged from t-tests and 0 otherwise. Those values were submitted to a 1D MCS. For each of the 10000 permutations, we randomized the binarized elements (threshold = .05) of the *p*-value time series and extracted the maximum cluster size. Then, we built a reference distribution of maximum cluster sizes and considered significant the original clusters that were larger than the 95% of the permuted ones.

## Author contributions

LB, EB, MLK and PV conceived the hypotheses and designed the study. LB, FC, JC, AS, DP, MLK performed pre-processing and statistical analysis. GD, MP, EB, MLK, DP, PCW and PV provided essential help to interpret and frame the results within the neuroscientific literature. LB wrote the first draft of the manuscript and, together with FC and MLK, prepared the figures. All the authors contributed to and approved the final version of the manuscript.

## Acknowledgements

We thank Giulia Donati, Riccardo Proietti, Giulio Carraturo, Mick Holt and Holger Friis for their assistance in the neuroscientific experiment. We also thank the psychologist Tina Birgitte Wisbech Carstensen for her help with the administration of psychological tests and questionnaires.

The Center for Music in the Brain (MIB) is funded by the Danish National Research Foundation (project number DNRF117). Additionally, we thank the Italian section *of Mensa: The International High IQ Society* for the economic support provided to the author Francesco Carlomagno and the University of Bologna for the economic support provided to the students Giulia Donati, Riccardo Proietti, Giulio Carraturo.

LB is supported by the Carlsberg Foundation and MIB.

MLK is supported by the ERC Consolidator Grant: CAREGIVING (n. 615539), Center for Music in the Brain, funded by the Danish National Research Foundation (DNRF117), and Centre for Eudaimonia and Human Flourishing funded by the Pettit and Carlsberg Foundations.

GD is supported by the Spanish Research Project PSI2016-75688-P (AEI/FEDER, EU), by the European Union’s Horizon 2020 Research and Innovation Programme under grant agreements n. 720270 (HBP SGA1) and n. 785907 (HBP SGA2), and by the Catalan AGAUR Programme 2017 SGR 1545.

JC is supported by Portuguese Foundation for Science and Technology CEECIND/03325/2017, Portugal.

## Competing interests statement

The authors declare no competing interests.

## SUPPLEMENTAL FIGURES

**Figure SF1.**
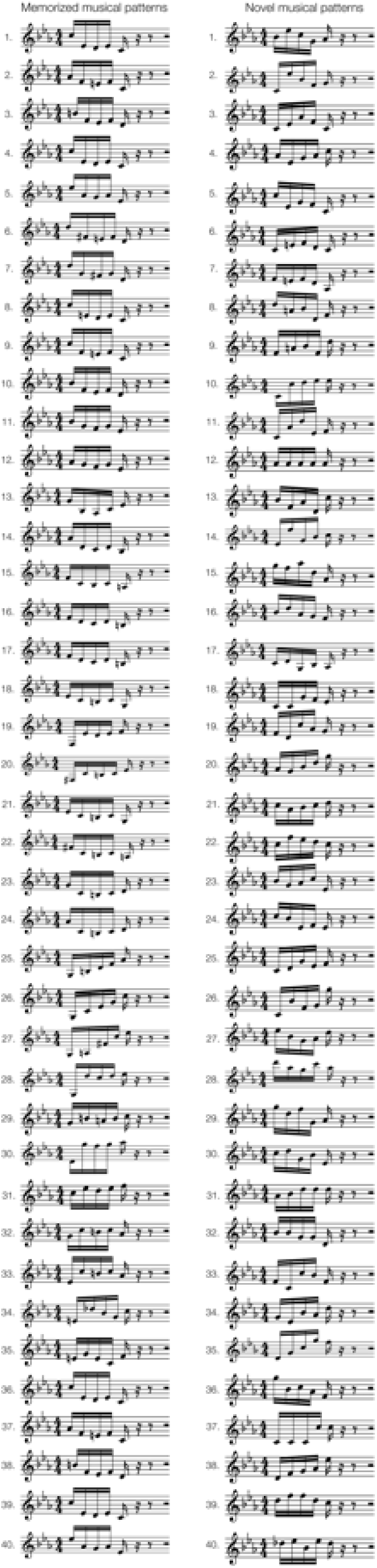
Musical patterns used in the experiment. Depiction of all musical patterns used in the auditory ‘old/new’ paradigm employed in the study. On the left, we have reported the 40 musical patterns extracted from Bach’s prelude (previously memorized musical patterns, ‘old’). On the right, we have reported the 40 novel melodies (novel musical patterns, ‘new’) that were created and matched to Bach’s prelude excerpts with regards to IC, H and main acoustic features.

**Figure SF2.**
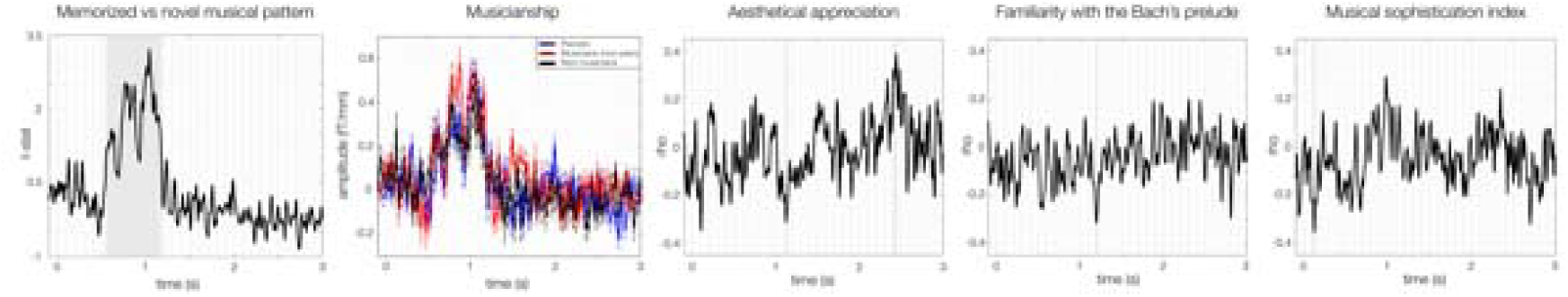
Brain activity underlying auditory pattern recognition and musical skills. The first plot from the left shows the contrast between brain activity underlying recognition of the previously memorized versus novel musical patterns. The second plot illustrates such contrast in relation to the musicianship groups of our sample (pianists, non-pianist musicians, non-musicians). The remaining plots show the correlation between the contrast and three measures of musical skills/features: aesthetical appreciation of the Bach’s prelude (i), familiarity with the Bach’s prelude (ii), general ability to be engaged with music and musical activities (iii).

**Figure SF3.**
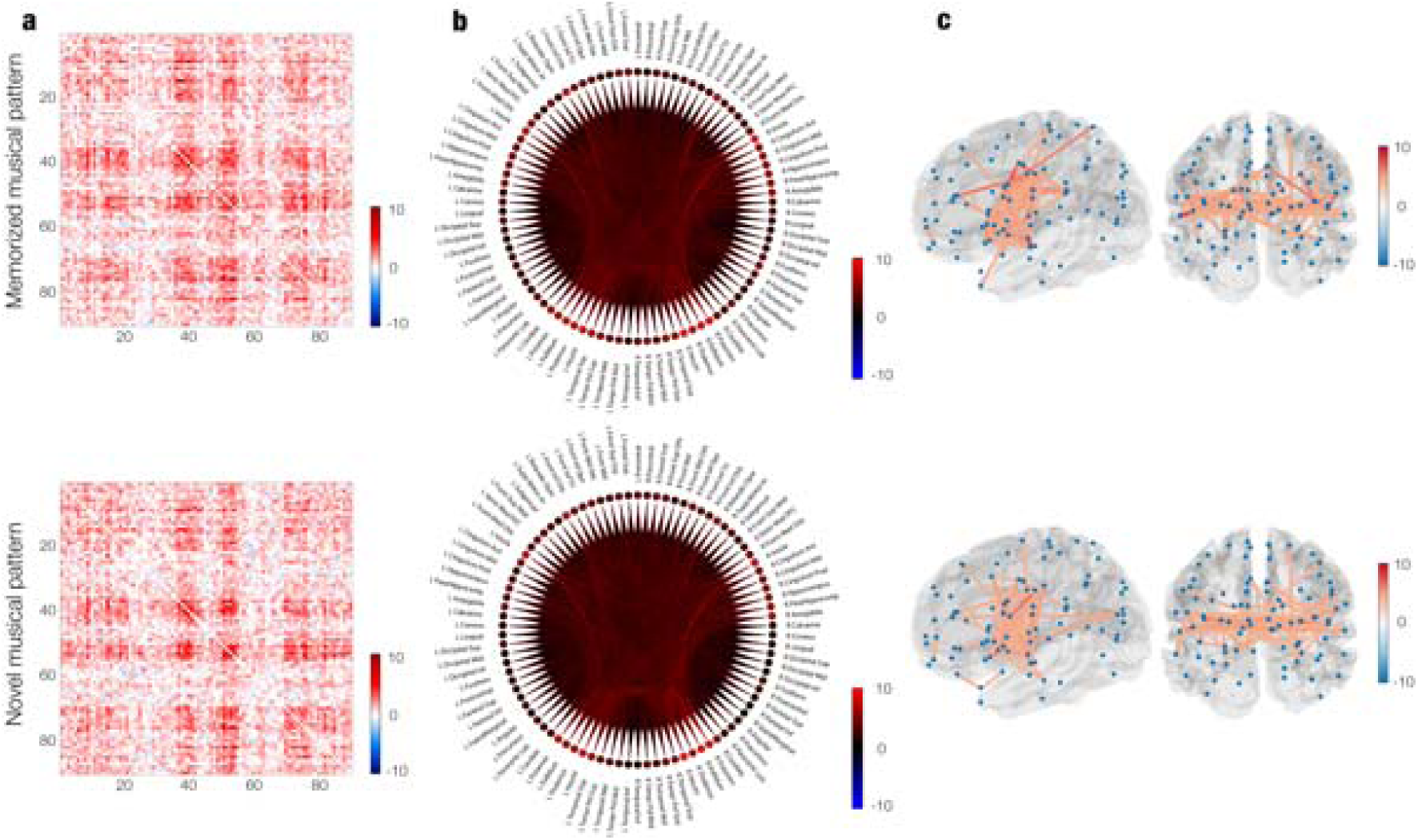
Static functional connectivity for theta band. **a –** Matrix representation of contrasts (t-values) between task versus baseline SFCs. **b –** Same values depicted in schemaballs. **c –** Same values depicted in brain templates. For each pair of brain templates, the left brain represents a left hemisphere perspective while the right one a posterior view of the brain.

**Figure SF4.**
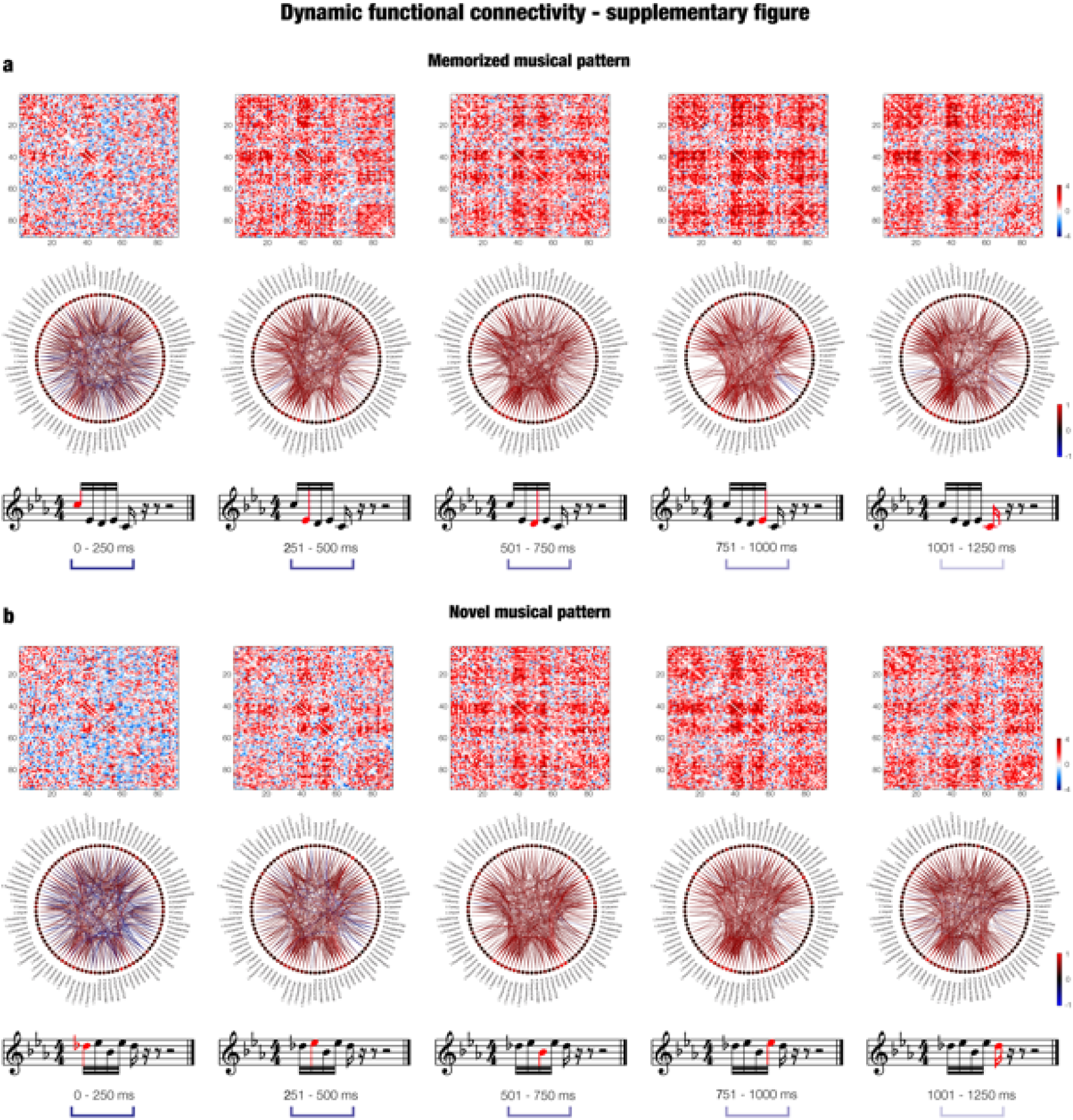
Dynamic functional connectivity II. **a –** DFC calculated for each tone composing the memorized musical patterns. The top row depicts the whole-brain connectivity in matrix representations, while the middle row in schemaball illustrations. Values refers to Wilcoxon sign-rank test z-values computed for memorized versus novel musical patterns DFC matrices. The bottom row illustrates an example of memorized musical pattern evolving over time. **b –** Same representation for novel musical patterns.

**Figure SF5.**
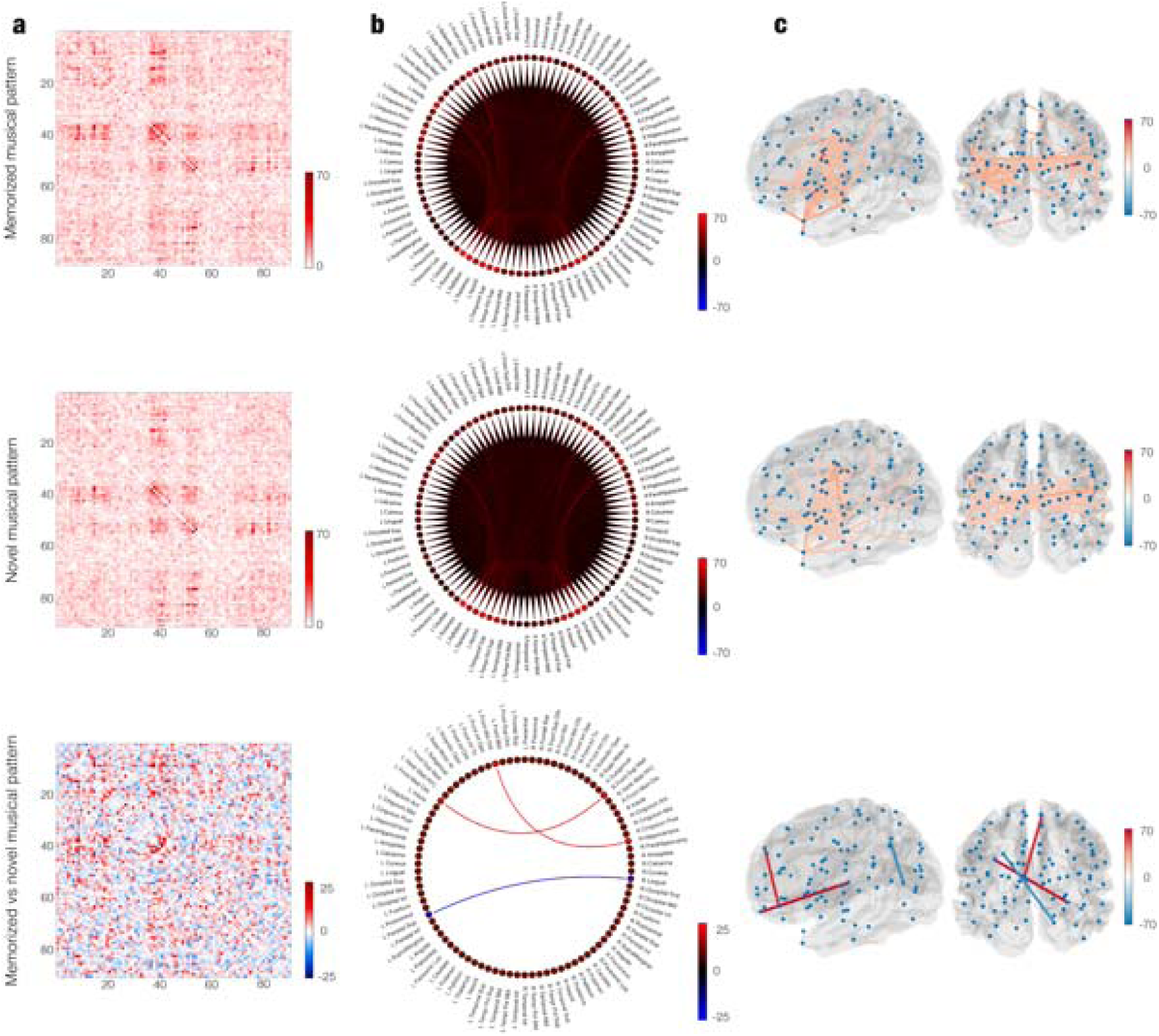
Phase synchrony coupling over time. **a –** Matrix representation of the sum over time (ms) of significant connections between each pair of brain areas contrasting memorized musical patterns versus baseline (top), novel musical patterns versus baseline (middle) and memorized versus novel musical patterns (bottom). b – Same results depicted in schemaballs. **c –** Same results illustrated in brain templates. For each pair of brain templates, the left brain represents a left hemisphere perspective while the right one a posterior view of the brain.

**Figure SF6.**
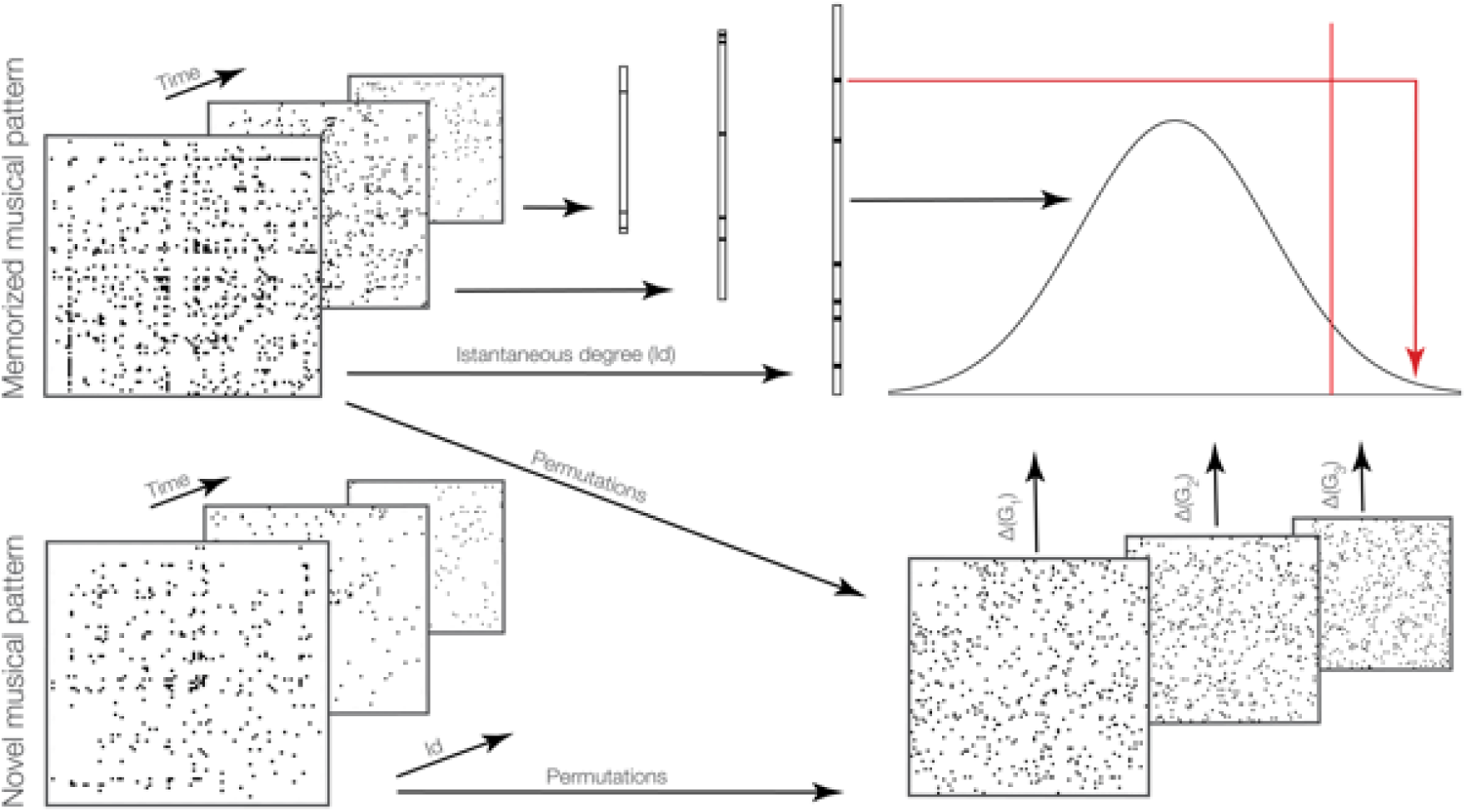
Example of degree centrality MCS algorithm. The Figure depicts the MCS algorithm for degree centrality of IFC matrices. Top left matrices reflect the IFC profile over time of the brain functional connectivity underlying recognition of the memorized and novel musical patterns. For each of those matrices, a number of permutations has been conducted shuffling their elements (bottom right). The resulting degrees of each matrix have been used to build the reference distribution (top right), here approximated by a normal distribution. Then, the significantly central ROIs within the whole-brain where the ones whose original degrees were higher than the X% of degrees obtained during the permutation: Here, X is a defined threshold described in the manuscript and slightly varying according to different implementations of MCS.

## SUPPLEMENTAL ITEM TITLES AND LEGENDS

### SUPPLEMENTARY TABLES

In the cases when the supplementary tables were too large to be conveniently reported in the current document, they have been reported in Excel files that can be found at the following link:

https://drive.google.com/drive/folders/1X3JAO-WRJ82roqDqcR9RhedenxhdfLgt?usp=sharing

**Table ST1.** Memorized versus novel musical patterns – MEG sensors

| **Cluster number** | **Size** | **Channels** | **Time-range (s)** | ***p*** |
| --- | --- | --- | --- | --- |
| *Gradiometers (memorized versus novel musical patterns)* |  |  |  |  |
| 1 | 2117 | 90 | 0. 547 – 1.180 | < .001 |
| *Magnetometers – positive (memorized versus novel musical patterns)* |  |  |  |  |
| 1 | 817 | 24 | 0.627 – 1.180 | < .001 |
| *Magnetometers – negative (memorized versus novel musical patterns)* |  |  |  |  |
| 1 | 190 | 18 | 0.727 – 0.880 | < .001 |
| 2 | 168 | 15 | 0.960 – 1.133 | < .001 |

**Table ST2.** Novel versus memorized musical patterns – MEG sensors

| Cluster number | Size | Channels | Time-range (s) | *p* |
| --- | --- | --- | --- | --- |
| *Gradiometers (novel versus memorized musical patterns)* |  |  |  |  |
| 1 | 14 | 2 | 1.640 – 1.713 | < .001 |
| 2 | 9 | 2 | 1.467 – 1.513 | < .001 |
| 3 | 8 | 2 | 2.387 – 2.427 | < .001 |
| 4 | 8 | 1 | 1.600 – 1.647 | < .001 |
| 5 | 8 | 2 | 1.760 – 1.793 | < .001 |
| 6 | 6 | 1 | 1.680 – 1.713 | < .001 |
| 7 | 6 | 2 | 0.487 – 0.507 | < .001 |
| 8 | 6 | 1 | 0.447 – 0.480 | < .001 |

**Table ST3 – Detailed information on significant clusters for MEG sensor data**

Significant clusters of MEG sensors emerged from MCS contrasting memorized versus novel musical patterns. The table depicts those clusters with regards to significant channels and time-windows.

**Table ST4 – Detailed information on significant clusters for MEG source data**

Significant clusters of MEG sources emerged from cluster-based permutation testing and related to memorized musical patterns versus baseline, novel musical patterns versus baseline and memorized versus novel musical patterns. The table depicts those clusters with regards to significant voxels, time-windows and averaged t-values for each voxel.

**Table ST5 – Brain activity for each tone of the musical excerpts**

Significant clusters of brain activity associated to each tone of the musical excerpts, reported for both conditions (memorized and novel musical patterns) and for their contrast (memorized versus novel musical patterns).

**Table ST6 – Brain areas significantly central within the whole-brain network for each musical tone and both experimental conditions**

Significantly central ROIs within the brain network over time, obtained contrasting independently memorized and novel musical patterns versus baseline.

**Table ST7.** Memorized versus novel musical patterns – ROIs centrality

| *Left central ROIs* | *p* | *Right central ROIs* | *p* |
| --- | --- | --- | --- |
| **Tone 1** |  |  |  |
| **Tone 2** |  |  |  |
| Middle frontal orbital cortex | < 1.0e-07 | Superior orbitofrontal cortex | < 1.0e-07 |
| Hippocampus | < 1.0e-07 | Middle cingulate gyrus | < 1.0e-07 |
| Middle frontal gyrus | < 1.0e-07 |  |  |
| Caudate | < 1.0e-07 |  |  |
| Fusiform gyrus | 8.8e-05 |  |  |
| **Tone 3** |  |  |  |
| Fusiform gyrus | < 1.0e-07 | Angular | < 1.0e-07 |
| Hippocampus | 2.2e-05 | Superior temporal gyrus | < 1.0e-07 |
|  |  | Ventro-med prefrontal cortex | 2.2e-05 |
|  |  | Fronto-superior medial cortex | 1.0e-04 |
| **Tone 4** |  |  |  |
| Caudate | < 1.0e-07 | Anterior cingulum gyrus | < 1.0e-07 |
| Precentral gyrus | 2.2e-05 | Inferior temporal gyrus | < 1.0e-07 |
|  |  | Fusiform gyrus | 1.1e-05 |
|  |  | Middle cingulum gyrus | 6.6e-05 |
| **Tone 5** |  |  |  |
| Parahippocampal gyrus | < 1.0e-07 | Parietal inferior lobule | < 1.0e-07 |
| Fronto-medial orbital cortex | < 1.0e-07 |  |  |
| Calcarine | < 1.0e-07 |  |  |
| Parietal superior lobule | < 1.0e-07 |  |  |
| Thalamus | < 1.0e-07 |  |  |
| Temporal pole middle | < 1.0e-07 |  |  |
| Rolandic operculum | 5.5e-05 |  |  |

**Table ST8.** Novel versus memorized musical patterns – ROIs centrality

| **Tone 1** |  |  |  |
| --- | --- | --- | --- |
| *Left central ROIs* | *p* | *Right central ROIs* | *p* |
| Precentral gyrus | < 1.0e-07 | Fronto-middle orbital cortex | 1.1e-05 |
| Fusiform gyrus | < 1.0e-07 | Supramarginal gyrus | 3.3e-05 |
| Cuneus | 7.8e-05 |  |  |
| Lingual gyrus | 3.3e-05 |  |  |
| **Tone 2** |  |  |  |
| Supramarginal gyrus | < 1.0e-07 | Middle frontal gyrus | < 1.0e-07 |
| Calcarine | 5.5e-05 | Occipital superior gyrus | < 1.0e-07 |
|  |  | Thalamus | < 1.0e-07 |
|  |  | Middle temporal gyrus | < 1.0e-07 |
|  |  | Lingual gyrus | 3.3e-05 |
| **Tone 3** |  |  |  |
| Middle frontal gyrus | < 1.0e-07 |  |  |
| Lingual | < 1.0e-07 |  |  |
| Supramarginal gyrus | < 1.0e-07 |  |  |
| Paracentral lobe | < 1.0e-07 |  |  |
| Cuneus | 7.7e-05 |  |  |
| Middle temporal gyrus | 2.2e-05 |  |  |
| **Tone 4** |  |  |  |
| Lingual gyrus | < 1.0e-07 | Precentral gyrus | < 1.0e-07 |
| Parietal inferior lobule | < 1.0e-07 | Supramarginal gyrus | < 1.0e-07 |
| Supramarginal gyrus | < 1.0e-07 | Middle frontal orbital cortex | < 1.0e-07 |
| Paracentral lobe | < 1.0e-07 | Ventro-med prefrontal cortex | 6.6e-05 |
| **Tone 5** |  |  |  |
| Occipital inferior gyrus | < 1.0e-07 | Fronto-middle orbital cortex | < 1.0e-07 |
| Post-central gyrus | 4.4e-05 | Inferior frontal orbital cortex | < 1.0e-07 |
|  |  | Inferior frontal gyrus, triangul | < 1.0e-07 |
|  |  | Superior frontal medial | < 1.0e-07 |
|  |  | Postcentral gyrus | < 1.0e-07 |
|  |  | Caudate | < 1.0e-07 |
|  |  | Middle temporal gyrus | < 1.0e-07 |

## SUPPLEMENTARY TEXT

### SR1 – Additional information on Monte-Carlo simulations on MEG combined gradiometers and magnetometers

We employed a different approach by calculating several univariate t-tests and then correcting for multiple comparisons by using MCS. Before computing the t-tests, in accordance with a large number of other MEG and electroencephalography (EEG) task studies [62], we averaged the trials over conditions, obtaining two mean trials, one for the memorized and one for the novel musical patterns. Then, we combined each pair of planar gradiometers by sum-root square. Afterwards, we computed a t-test for each combined planar gradiometer and each time-point in the time-range 0 – 2.500 seconds, contrasting the two experimental conditions. We reshaped the matrix for obtaining, for each time-point, a 2D approximation of the MEG channels layout that we binarized according to the *p*-values obtained from the previous t-tests (threshold = .01) and the sign of *t*-values. The resulting 3D matrix (*M*) was therefore composed by 0s when the t-test was not significant and 1s when it was. Then, to correct for multiple comparisons, we identified the clusters of 1s and assessed their significance by running MCS. Specifically, we made 1000 permutations of the elements of the original binary matrix *M*, identified the maximum cluster size of 1s and built the distribution of the 1000 maximum cluster sizes. Finally, we considered significant the original clusters that had a size bigger than the 99.9% maximum cluster sizes of the permuted data. Considering that magnetometers (differently from combined gradiometers) maintain the double polarity of the magnetic field, contrasting two experimental conditions presents potential technical ambiguities, admitting the theoretical possibility that two neighboring clusters with opposite polarity and depicting different strengths between conditions (e.g. cond1 > cond2 (positive polarity) in cluster one and cond2 > cond1 (negative polarity) in cluster two) may be identified as one unique large (positive) cluster. For these reasons, at first, we carried out the algorithm by contrasting memorized versus novel musical patterns for combined planar gradiometers only. Then, on the basis of the significant clusters emerged, we used the same algorithm one more time for magnetometers only, within the significant time-range emerging from the first MCS (in this case: 0.547 – 1.180 seconds). This procedure allowed us to obtain more reliable and complete information about the different neural signal associated to the recognition of the memorized and novel musical patterns for both gradiometers and magnetometers. The whole MCS procedure was performed for memorized versus novel musical patterns and vice versa.

### SR2 - Significantly central ROIs within the whole-brain network detected from the SFC matrices for delta, alpha, beta and gamma bands

We observed left cingulum middle (*p* < 1.0e-07), parahippocampal gyrus (*p* = 5.5e-06), amygdala (*p* = 2.2e-06), Heschl’s gyrus (*p* = 5.5e-06), post-central gyrus (*p* = 1.0e-04), thalamus (*p* < 1.0e-07), right thalamus (*p* = 2.2e-06), pallidum (*p* < 1.0e-07), putamen (*p* < 1.0e-07), cuneus (*p* < 1.0e-07), caudate (*p* = 3.3e-06), cingulum anterior (*p* < 1.0e-07) and middle (*p* = 7.7e-05), insula (*p* < 1.0e-07) for delta; left precentral gyrus (*p* = 8.8e-05), cingulum middle (*p* < 1.0e-07) and posterior (*p* = 1.1e-06), hippocampus (*p* < 1.0e-07), Heschl’s gyrus (*p* < 1.0e-07), right superior temporal gyrus (*p* < 1.0e-07), thalamus (*p* = 6.6e- 05), Heschl’s gyrus (*p* < 1.0e-07), caudate (*p* = 5.0e-05), cingulum middle (*p* = 2.6e-05), subgenual cortex (*p* < 1.0e-07), Rolandic operculum (*p* < 1.0e-07), fronto-superior orbital cortex (*p* = 1.5e-05), fronto-medial orbital cortex (*p* = 1.8e-05) for alpha; left supplementary motor area (*p* = 5.6e-05), cingulum middle (*p* < 1.0e-07), caudate (*p* = 2.2e-06), right temporal pole superior (*p* = 1.1e-06), Heschl’s gyrus (*p* = 1.1e-06), thalamus (*p* < 1.0e-07), putamen (*p* = 2.2e-06), caudate (*p* < 1.0e-07), pallidum (*p* = 4.4e-06), parahippocampal gyrus (*p* = 4.8e-06), cingulum posterior (*p* < 1.0e-07), middle (*p* < 1.0e-07) and anterior (*p* < 1.0e- 07), amygdala (*p* = 1.1e-05), fronto-medial orbital cortex (*p* < 1.0e-07), subgenual cortex (*p* < 1.0e-07) for beta; left cingulum anterior (*p* < 1.0e-07) and middle (*p* < 1.0e-07), caudate (*p* < 1.0e-07), putamen (*p* < 1.0e-07), pallidum (*p* < 1.0e-07), right pallidum (*p* < 1.0e-07), putamen (*p* < 1.0e-07), caudate (*p* < 1.0e-07), cingulum anterior (*p* < 1.0e-07), subgenual cortex (*p* < 1.0e-07), fronto-medial orbital cortex (*p* < 1.0e-07), precentral gyrus (*p* < 2.2e-06) for gamma.

These results are illustrated in **Figure 5**.

### SR3 – Whole-brain phase synchronization coupling

We also computed the phase synchronization coupling (PSC) of brain areas *n_j_* and *n_m_* over the entire task, according to the following equation:

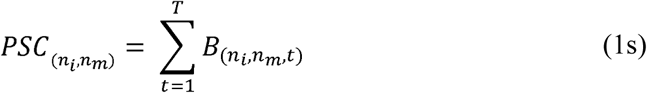

We obtained two symmetric square matrices, one for the memorized musical patterns and one for the novel ones, by summing over time the IFC matrices of both conditions. We used a further MCS to assess the significance of the differences between conditions for each pair of brain areas (summarized in the matrix *CD)*. We made 10000 permutations of the upper triangles of the memorized and novel musical pattern matrices and we calculated the difference matrix *CDP_p_* for each permutation. Then we created two distributions, for the maximum and minimum values of *CDP_p_* for each permutation. Finally, we compared the positive values of *CD* with the distribution of the maximums and the negative values of *CD* with the distribution of the minimums. Considering that we hypothesized to detect significant differences between the two experimental conditions among some brain connections but without specifying which ones, we had to correct the final results taking into account the total number of brain connections (4005 since they are undirected) and the directions of the contrasts (two). Therefore, the threshold for considering a couple of brain areas significant was set as: 6.2e-05 (0.05/(4005*2)). According to this threshold, we detected the two following significant connections that occurred more frequently for memorized versus novel musical patterns: left middle frontal orbital cortex and right hippocampus appeared significant (*p <* 6.2e-05); left frontal medial orbital cortex and right frontal superior medial cortex (*p <* 6.2e-05). Conversely, novel versus memorized musical patterns returned the following significant connection between brain regions: left parietal superior lobule and right lingual gyrus (*p <* 6.2e-05). The results for memorized, novel musical patterns, and their contrast are depicted in **Figure SF5**.

## Notes

### Competing Interest Statement

The authors have declared no competing interest.

## References

[1] Friston, K., & Kiebel, S. Predictive coding under the free-energy principle. Philosophical Transactions of the Royal Society B: Biol Scien, 364(1521), 1211–1221 (2009).

[2] Koelsch, S., Vuust, P., & Friston, K. Predictive Processes and the Peculiar Case of Music. Tren in Cog Scien, 23(1), 63–77 (2019).

[3] Dehaene, S., Kerszberg, M., & Changeux, J. P. A neuronal model of a global workspace in effortful cognitive tasks. Proc of the Nat Aca of Scien of the Unit Stat of Ame, 95(**24**), 14529–14534 (1998).

[4] Patel, A. D. Music, Language, and the Brain (Oxford University Press, Oxford, 2008).

[5] Wixted, J. T. Dual-process theory and signal-detection theory of recognition memory. Psychol Rev 114(**1**), 152–176 (2007).

[6] Yonelinas, A. P. The nature of recollection and familiarity: A review of 30 years of research. Jour of Mem and Lang 46(**3**), 441–517 (2002).

[7] Eichenbaum, H., Yonelinas, A. P., & Ranganath, C. The Medial Temporal Lobe and Recognition Memory. Ann Rev of Neuroscien, 30, 123–152 (2007).

[8] Aggleton, J. P., & Brown, M. W. Interleaving brain systems for episodic and recognition memory. Trends in Cog Scien, 10(**10**), 455–643. (2006).

[9] DiCarlo, J. J., Zoccolan, D., & Rust, N. C. How does the brain solve visual object recognition? Neuron, 73(3), 415–34 (2012).

[10] Brattico, E., Näätänen, R., & Tervaniemi, M. Context effects on pitch perception in musicians and nonmusicians: Evidence from event-related potential recordings. Mus Per, 19(**2**), 199–222 (2001).

[11] Näätänen, R., Paavilainen, P., Rinne, T., & Alho, K. The mismatch negativity (MMN) in basic research of central auditory processing: A review. Clin Neurophy 118(**12**), 2544–2590 (2007).

[12] Dehaene, S., Meyniel, F., Wacongne, C., Wang, L., & Pallier, C. The Neural Representation of Sequences: From Transition Probabilities to Algebraic Patterns and Linguistic Trees. Neuron, 88(**1**), 2–19 (2015).

[13] Peretz, I., & Zatorre, R. J. The Cognitive Neuroscience of Music (Oxford University Press, Oxford, 2003).

[14] Raynor, H., & Cooke, D. The Language of Music (Oxford University Press, Oxford, 1960).

[15] Jackendoff R. Parallels and nonparallels between language and music. Mus Perc, 26(3), 195–204 (2009).

[16] Plack, C. J. The sense of hearing (Lawrence Erlbaum Associates Publishers, Mahwah, 2005).

[17] Williams, P. J. S. Bach: A Life in Music (Cambridge University Press, Cambridge, 2007).

[18] Hofstadter, D. R. Gödel, Escher, Bach: An Eternal Golden Braid (Basic Books, New York, 1999).

[19] King, J.R. and Dehaene, S. Characterizing the dynamics of mental representations: the temporal generalization method. Tren in Cog Scien, 18(*4*), 203–210 (2014).

[20] Salti, M., Monto, S., Charles, L., King, J.R., Parkkonen, L. and Dehaene, S. Distinct cortical codes and temporal dynamics for conscious and unconscious percepts. Elife, 4, p.e05652 (2015).

[21] Changeux, J. P. Secrets of Creativity: What Neuroscience, the Arts, and Our Minds Reveal. (Oxford University Press, Oxford, 2019).

[22] Stark, E. A., Vuust, P., & Kringelbach, M. L. Music, dance, and other art forms: New insights into the links between hedonia (pleasure) and eudaimonia (well-being). Prog in Brain Res 237, 129–152 (2018).

[23] Näätänen, R., Pakarinen, S., Rinne, T., & Takegata, R. The mismatch negativity (MMN): Towards the optimal paradigm. Clin Neurophy 115(**1**), 140–144 (2004).

[24] Remedios, R., Logothetis, N. K., & Kayser, C. An auditory region in the primate insular cortex responding preferentially to vocal communication sounds. Journ of Neuroscien 29(**4**), 1034–1045 (2009).

[25] Bird, C. M. The role of the hippocampus in recognition memory. Cortex 93, 155–165 (2017).

[26] Brown, M. W., & Aggleton, J. P. Recognition memory: What are the roles of the perirhinal cortex and hippocampus? Nat Rev Neuroscien, 2(**1**), 51–61 (2001).

[27] Teixeira, C. M., Pomedli, S. R., Maei, H. R., Kee, N., & Frankland, P. W. Involvement of the anterior cingulate cortex in the expression of remote spatial memory. Journ of Neuroscien 26(**29**), 7555–7564 (2006).

[28] Bach, D. R. et al. The effect of appraisal level on processing of emotional prosody in meaningless speech. NeuroImage, 42(**2**), 919–927 (2008).

[29] Stephenson-Jones et al. A basal ganglia circuit for evaluating action outcomes. Nature 539(7628), 289–293 (2016).

[30] Kringelbach, M. L. The hedonic brain: a functional neuroanatomy of human pleasure. Pleasures of the Brain, 13(11), 479–487 (2009).

[31] Vuust, P., & Kringelbach, M. L. The pleasure of making sense of music. Int Scien Rev 35(**2**), 166–182 (2010).

[32] Zatorre, R. J., Belin, P., & Penhune, V. B. Structure and function of auditory cortex: Music and speech. Tren in Cog Scien 6(**1**), 37–46 (2002).

[33] Limongi, R., Sutherland, S. C., Zhu, J., Young, M. E., & Habib, R. Temporal prediction errors modulate cingulate-insular coupling. NeuroImage, 71, 147–157 (2013).

[34] Koelsch, S., Fritz, T., Cramon, D. Y. V., Müller, K., & Friederici, A. D. Investigating emotion with music: An fMRI study. Hum Brain Map, 27(3), 239–250 (2006).

[35] Stephenson-Jones, M., Kardamakis, A. A., Robertson, B., & Grillner, S. Independent circuits in the basal ganglia for the evaluation and selection of actions. Proc of the Nat Aca of Scien of the Unit Stat of Ame a 110(**38**), 3670–3679 (2013).

[36] Squire, L. R., & Bayley, P. J. The neuroscience of remote memory. Cur Opin in Neurob 17(**2**), 185–196 (2007).

[37] Packard, M. G., & Knowlton, B. J. Learning and Memory Functions of the Basal Ganglia. Ann Rev of Neuroscien 25, 563–593 (2002).

[38] Dehaene, S., & Changeux, J. P. Experimental and Theoretical Approaches to Conscious Processing. Neuron, 70(**2**), 200–227 (2011).

[39] Dehaene, S., Changeux, J. P., & Naccache, L. The global neuronal workspace model of conscious access: From neuronal architectures to clinical applications. Res and Persp in Neuroscien, 55–84 (2011).

[40] Dehaene, S., & Changeux, J. P. Ongoing spontaneous activity controls access to consciousness: A neuronal model for inattentional blindness. PLoS Biol, 3(**5**), e141 (2005).

[41] Brown, J. W., & Braver, T. S. Learned predictions of error likelihood in the anterior cingulate cortex. Science, 307(5712), 1118–1121 (2005).

[42] Holroyd, C. B. et al. Dorsal anterior cingulate cortex shows fMRI response to internal and external error signals. Nat Neuroscien, 7(5), 497–498 (2004).

[43] Mack, M. L., Love, B. C., & Preston, A. R. Building concepts one episode at a time: The hippocampus and concept formation. Neuroscien Let 680, 31–38 (2018).

[44] Shohamy, D., & Wagner, A. D. Integrating Memories in the Human Brain: Hippocampal-Midbrain Encoding of Overlapping Events. Neuron 60(**2**), 378–389 (2008).

[45] Näätänen, R., & Picton, T. The N1 Wave of the Human Electric and Magnetic Response to Sound: A Review and an Analysis of the Component Structure. Psychophy 24(**4**), 375–425 (1987).

[46] Parras, G. et al. Neurons along the auditory pathway exhibit a hierarchical organization of prediction error. Nat Com 8(**1**), 1–17(2017).

[47] Wacongne, C. et al. Evidence for a hierarchy of predictions and prediction errors in human cortex. Proc of the Nat Aca of Scien of the Unit Stat of Ame 108 (**51**), 20754–20759 (2011).

[48] Näätänen, R., Tervaniemi, M., Sussman, E., Paavilainen, P., & Winkler, I. “Primitive intelligence” in the auditory cortex. Tren in Neuroscien 24(**5**), 283–288 (2001).

[49] Walter, W. G., Cooper, R., Aldridge, V. J., McCallum, W. C., & Winter, A. L. Contingent negative variation: An electric sign of sensori-motor association and expectancy in the human brain. Nature 203, 380–384 (1964).

[50] Bonetti, L., Haumann, N.T., Brattico, E., Kliuchko, M., Vuust, P., Särkämö, T. and Näätänen, R. Auditory sensory memory and working memory skills: Association between frontal MMN and performance scores. Brain res, 1700, 86–98 (2018).

[51] Kayser, J., Fong, R., Tenke, C. E., & Bruder, G. E. Event-related brain potentials during auditory and visual word recognition memory tasks. Cogn Brain Res, 16(*1*), 11–25 (2003).

[52] Pearce, M. T. Statistical learning and probabilistic prediction in music cognition: Mechanisms of stylistic enculturation. Ann of the New York Aca of Scien 1423(1), (2018).

[53] Taulu, S. & Simola, J. Spatiotemporal signal space separation method for rejecting nearby interference in MEG measurements. Phys in Med and Biol 51, 1759–1768 (2006).

[54] Woolrich, M. W., Hunt, L., Groves, A., Barnes, G. MEG beamforming using Bayesian PCA for adaptive data covariance matrix regularization. NeuroImage 57(**4**), 1466–1479 (2011).

[55] Woolrich, M. W. et al. Bayesian analysis of neuroimaging data in FSL. NeuroImage 45, S173–S186 (2009).

[56] Penny, W., Friston, K., Ashburner, J., Kiebel, S., & Nichols, T. Statistical Parametric Mapping: The Analysis of Functional Brain Images. Statistical Parametric Mapping: The Analysis of Functional Brain Images. (Academic Press, Cambridge, USA, 2007).

[57] Oostenveld, R., Fries, P., Maris, E., Schoffelen, J. M. FieldTrip: Open source software for advanced analysis of MEG, EEG, and invasive electrophysiological data. Compl Int and Neuroscien 2011, 156869 (2011).

[58] Mantini, D. et al. A signal-processing pipeline for magnetoencephalography resting-state networks. Brain Con 1(**1**), 49–59 (2011).

[59] Wilson, M. D. Support Vector Machines. In Encyclopedia of Ecology, Five-Volume Set, (Academic Press, Cambridge, USA, 3431–3437, 2008).

[60] Haufe, S. et al. On the Interpretation of Weight Vectors of Linear Models in Multivariate. Neuroimaging 87, 96–110 (2014).

[61] Cichy, R. M., Pantazis, D., & Oliva, A. Resolving human object recognition in space and time. Nat Neuroscien, 17(**3**), 455–462 (2014).

[62] Gross, J. et al. Good practice for conducting and reporting MEG research. NeuroImage 65, 349–363 (2013).

[63] Müllensiefen, Daniel, Bruno Gingras, Jason Musil, and Lauren Stewart. The musicality of non-musicians: an index for assessing musical sophistication in the general population. PloS one 9, *2*. (2014). e89642.

[64] Wechsler, D. Wechsler Memory Scale. The Psychological Corporation (1997).

[65] Hillebrand, A. & Barnes, G. R. Beamformer analysis of MEG data. International Review of Neurobiology 68, 149–171 (2005).

[66] Brookes, M. J. et al. Beamformer reconstruction of correlated sources using a modified source model. NeuroImage 34, 1454–1465 (2007).

[67] Hunt, L. T. et al. Mechanisms underlying cortical activity during value-guided choice. Nature Neuroscience 15, 470–476 (2012).

[68] Tzourio-Mazoyer, N. et al. Automated anatomical labeling of activations in SPM using a macroscopic anatomical parcellation of the MNI MRI-single-subject brain. NeuroImage 15, 273–289 (2002).

[69] Brookes, M. J. et al. A multi-layer network approach to MEG connectivity analysis. NeuroImage 132, 425– 438 (2016).

[70] Colclough, G. L., Brookes, M. J., Smith, S. M. & Woolrich, M.W. A symmetric multivariate leakage correction for MEG connectomes. NeuroImage 117, 439–448 (2015).

[71] Lee, D. J., Kulubya, E., Goldin, P., Goodarzi, A., & Girgis, F. Review of the neural oscillations underlying meditation. Front in Neuroscien 12, 178 (2018).

[72] Allen, M., Poggiali, D., Whitaker, K., Marshall, T. R., Kievit, A. R. Raincloud Plots: A Multi-Platform Tool for Robust Data Visualization. Well Open Res 1, 4:63 (2019).

[73] Kroese, D. P., Taimre, T., & Botev, Z. I. Handbook of Monte Carlo Methods. Handbook of Monte Carlo Methods (John Wiley and Sons, New York, 2011).

[74] Rubinov, M. & Sporns, O. Complex network measures of brain connectivity: Uses and Interpretations. NeuroImage 52, 1059–1069 (2010).

[75] Cabral, J. et al. Cognitive performance in healthy older adults relates to spontaneous switching between states of functional connectivity during rest. Scien Rep 7, 5135 (2017).

[76] Kuramoto, Y. Chemical Oscillations, Waves, and Turbulence (Springer-Verlag, New York, 1984).

